# Divergent ecological responses to typhoon disturbance revealed via landscape-scale acoustic monitoring

**DOI:** 10.1101/2023.01.11.523682

**Authors:** Samuel R.P-J. Ross, Nicholas R. Friedman, Kenneth L. Dudley, Takuma Yoshida, Masashi Yoshimura, Evan P. Economo, David W. Armitage, Ian Donohue

## Abstract

Climate change is increasing the frequency, intensity, and duration of extreme weather events across the globe. Understanding the capacity for ecological communities to withstand and recover from such events is critical. Typhoons are extreme weather events that are expected to broadly homogenise ecological dynamics through structural damage to vegetation and longer-term effects of salinization. Given their unpredictable nature, monitoring ecological responses to typhoons is challenging, particularly for mobile animals such as birds. Here, we report spatially variable ecological responses to typhoons across terrestrial landscapes. Using a high temporal resolution passive acoustic monitoring network across 24 sites on the subtropical island of Okinawa, Japan, we found that typhoons elicit divergent ecological responses among Okinawa’s diverse terrestrial habitats, as indicated by increased spatial variability of biological sound production (biophony) and individual species detections. This suggests that soniferous communities are capable of a diversity of different responses to typhoons. That is, spatial insurance effects among local ecological communities provide resilience to typhoons at the landscape scale. Even though site-level typhoon impacts on soundscapes and bird detections were not particularly strong, monitoring at scale with high temporal resolution across a broad spatial extent nevertheless enabled detection of spatial heterogeneity in typhoon responses. Further, species-level responses mirrored those of acoustic indices, underscoring the utility of such indices for revealing insight into fundamental questions concerning disturbance and stability. Our findings demonstrate the significant potential of landscape-scale acoustic sensor networks to capture the understudied ecological impacts of unpredictable extreme weather events.

## Introduction

Climate change is increasing both the frequency and destructive potential of extreme weather events such as typhoons (Bhatia et al., 2019; Emanuel, 2005; Kossin et al., 2020; Li & Chakraborty, 2020). Typhoons bring exceptional winds and rainfall and can cause structural damage to terrestrial habitats through wind-damage (Everham & Brokaw, 1996), flooding (Gardner et al., 1991), and salinization by sea spray, particularly on islands (Elliott & Nino, 1960; Kerr, 2000). These destructive forces can impact the mortality, composition, and dynamics of trees (Lin et al., 2020; Morimoto et al., 2021), birds (Cely, 1991; Chevalier et al., 2019; Seki, 2005), bacteria (Ares et al., 2020), invertebrates (Azuma et al., 1997; Willig & Camilo, 1991), and other animals (Donihue et al., 2018; Pavelka et al., 2007; Testard et al., 2021).

Land cover may affect exposure to—and hence the ecological consequences of—extreme weather events (Laurance, 1998). Several studies suggest that primary forest with a mix of vegetation types and life forms is most likely to resist and recover from typhoon disturbance (Abbas et al., 2020; Zampieri et al., 2020). Human activity and associated land use change has, therefore, considerable potential to modify the severity and reach of typhoons (Raymond et al., 2020). For example, deforestation through urbanisation or agricultural intensification leads to drastically simplified canopy structure, thereby increasing typhoon exposure, as well as subjecting ecological communities to the stressors and pollutants associated with these anthropogenic land uses (Daskalova et al., 2020; Senzaki et al., 2020; Uchida & Ushimaru, 2014). In contrast, land abandonment might be expected to dampen the effects of typhoons over time as intensively managed agricultural areas undergo natural succession to a more biologically and structurally diverse system (Sirami et al., 2008; Uchida & Ushimaru, 2014). Yet, despite the potential for land cover to moderate the impact of extreme events, the practical difficulties associated with ecological monitoring at scale have, to date, limited understanding of the extent of any differential impacts of typhoons across habitat types and of the consequences of such differences for landscape-scale biodiversity or spatial processes such as metacommunity dynamics and stability (Loreau et al., 2003; Wang et al., 2021).

Ecological stability is a central framework for considering disturbance impacts across spatial, temporal, and organisational scales, from populations to ecosystems (Hillebrand et al., 2018; Kéfi et al., 2019). Stability is a concept with multiple dimensions (Donohue et al., 2013; Hillebrand et al., 2018; Pimm, 1984), including components such as resistance to and recovery from disturbance (Baert et al., 2016; Yang et al., 2019), and the variability of ecological variables both in time and space (Tilman et al., 2006; Wang et al., 2017). Disturbance events and ecological responses to such events vary across spatiotemporal scales (Clark et al., 2021; Ross et al., 2021b; Zelnik et al., 2018). This necessitates high-resolution and long-term monitoring of ecosystems to holistically capture the ecological impacts of infrequent extreme events such as typhoons. However, monitoring biodiversity over large spatial and temporal scales poses considerable logistical and financial challenges. Accordingly, most empirical studies of ecological stability are experimental (Kéfi et al., 2019), while observational studies of disturbance typically employ space-for-time substitutions (Butsic et al., 2017), or consider only single-time snapshots before and after disturbance (*e.g.*, Burivalova et al., 2014). In such cases, it is extremely challenging to isolate the relevant pathways through which disturbance events impact ecosystems in a holistic multidimensional way.

Recent advances in automation hold promise for understanding disturbance responses through large-scale continuous monitoring of biodiversity (Keitt & Abelson, 2021; Ross et al., 2023). Following developments in data acquisition, storage, and processing, passive acoustic monitoring of wildlife and soundscapes is growing in popularity (Burivalova et al., 2019; Gibb et al., 2019). As sensor networks are established to collect acoustic data autonomously (Keitt & Abelson, 2021; Sethi et al., 2020), a diverse range of ecological studies become tractable by leveraging high-resolution acoustic time series (*e.g.*, Deichmann et al., 2018; Lomolino et al., 2015; Rossi et al., 2017; Sueur et al., 2019; Ross et al., 2023). Studies of disturbance impacts on *soundscapes*—that is, all sound produced in an ecosystem (Pijanowski et al., 2011a, 2011b), including *biophony* (biotic sound), *geophony* (natural abiotic sound, such as rain), and *anthropophony* (human-related sound)—have recently emerged, though, as with traditional studies of disturbance responses, most acoustic studies make before-and-after or space-for-time comparisons to assess disturbance impacts (*e.g.*, Deichmann et al., 2017; Gasc et al., 2018). However, the high-resolution time series afforded by passive acoustic monitoring allows opportunistic measurement of soundscape responses to infrequent disturbance events, such as typhoons (*e.g.*, Gottesman et al., 2021), as well as documenting longer-term trends under climate change (Sueur et al., 2019). Acoustic monitoring thus provides an opportunity to overcome many of the challenges associated with studying extreme weather events, by allowing pre- and post-typhoon comparisons (Altwegg et al., 2017; Rajan et al., 2022), and capturing ecological responses to typhoons across scales in space and time (Lin et al., 2020) using a multidimensional stability framework (Donohue et al., 2013). Of the few studies that have used acoustic monitoring to capture storms or extreme events, most focused on marine soundscapes (Boyd et al., 2021; Locascio & Mann, 2005; Simmons et al., 2021), though Gottesman *et al*. (2021) recently showed that terrestrial soundscapes were less resistant than those of coral reefs to hurricane disturbance. Embedded within terrestrial soundscapes, bird vocalisations provide the opportunity to assess the impact of typhoons on critical indicator taxa (Gasc et al., 2017), while acoustic indices provide rapid information on a combination of biodiversity and other meaningful aspects of soundscape change (Bradfer-Lawrence et al., 2020; Harris et al., 2016; Rajan et al., 2022; Müller et al., 2023; Sethi et al., 2023). There are, however, few studies that simultaneously assess both individual species vocalisations and acoustic indices explicitly (Ferreira et al., 2018; Ross et al., 2018).

Here, we exploit a dataset that is, to our knowledge, the highest-resolution dataset recording biological responses to an extreme weather event to date, capturing daily bird vocalisations and half-hourly acoustic indices in response to two large typhoons across 24 field sites on the island of Okinawa, Japan. We measure multiple dimensions of ecological stability for both soundscapes and individual bird species in response to a super-typhoon in September 2018, which was followed five days later by an extratropical cyclone. Our study spans Okinawa’s full range of terrestrial habitats, allowing us to examine how land use can shape ecological responses to extreme events. Given that organisms are differently vulnerable to mortality and mechanical damage resulting from typhoons (Abbas et al., 2020; Zampieri et al., 2020), we expect land use to influence typhoon responses (Raymond et al., 2020). We focus here on immediate and short-term responses of soundscapes to typhoon disturbance, with the aim of revealing the homogenising effects of typhoons on soundscapes and vocalising animal communities across the Okinawan landscape.

Specifically, we test the hypotheses that typhoons (1) temporarily reduce soundscape richness and (2) bird vocalisation rates, and (3) homogenise soundscapes across sites. We also predict that (4) natural forest habitats should have soundscapes that are more resistant to typhoons owing to their closed canopy structure (Abbas et al., 2020; Nimmo et al., 2016). We expect to find differences in bird species responses to typhoons, perhaps as a function of their traits (Wiley & Wunderle, 1993). Closed canopy specialists, frugivores, granivores, and nectarivores should be most vulnerable to food resource losses following typhoons (Chevalier et al., 2019; Wiley & Wunderle, 1993; Zhang et al., 2016), while insectivores may benefit from increased access to prey in canopy gaps (Cely, 1991; Seki, 2005), and large-bodied or predatory birds may be especially vulnerable to typhoon-induced habitat alteration, owing to their dependence on prey availability and habitat area (*e.g.*, Ross et al., 2019) and their slow reproductive rates (Cely, 1991; Cohen et al., 2021; Wiley & Wunderle, 1993; see Table S1a for summaries of our hypotheses for the focal species in this study). These hypotheses draw on the idea that forest loss is a key catalyst of biodiversity change (Daskalova et al., 2020; Gibson et al., 2011). This is especially pertinent given the high richness and rates of endemicity and specialism among Okinawa’s forest taxa (Inoue et al., 2019; Itô et al., 2000). Okinawa island is subject to rapid and ongoing land use change, particularly through deforestation for urbanisation and agricultural intensification (Ross et al., 2018; Takeuchi et al., 1981). Such land use change necessitates an explicit focus on habitat degradation as a driver of the ecological outcomes of intensifying natural disturbance regimes under climate change, including an increase in the frequency and destructive potential of typhoons and extreme storms around Okinawa (A. Iwasaki, unpublished data).

## Methods

### Study sites and typhoon impact

This study uses data from the OKEON (Okinawa Environmental Observation Network) Churamori Project (OKEON 美ら森プロジェクト; https://okeon.unit.oist.jp/) in Okinawa, Japan. We use data from OKEON’s 24 field sites across the island of Okinawa, representing Okinawa’s full range of land cover types (Figure 1; elevational range: 0.65 – 374.5 m). Elsewhere, we describe the geographic variation among the sites (Ross *et al*., 2018), which were assessed using reflectance estimates from Landsat 8 images to estimate proportional land cover for various land cover classes within a 1,000 m circular buffer surrounding each site. This buffer size was chosen as it represents a trade-off between local habitat structure and regional landscape features, which together shape the diversity of highly mobile taxa such as birds (Clough *et al*., 2009). Three pairs of sites had partly overlapping buffer areas at this scale: Kurashiki and Tounan (6.5% overlap), OIST forest and OIST campus (21% overlap), and Senbaru and Uehara (60% overlap). The centroid of all sites was sufficiently distant such that no individual animals could be simultaneously detected by more than a single recording unit, so we deemed our measurements from these sites to be suitably independent for this study. We classified land cover into the following categories: dense closed-canopy forest; grassland and scrubland (that is, low-medium growth coastal and disturbed vegetation, and managed grasses); agricultural land (primarily for sugarcane); urban areas characterised by materials such as asphalt and concrete with limited vegetation; sand and dirt with limited vegetation; freshwater bodies; and miscellaneous land cover not described in the above categories. To deal with the challenge of multicollinearity among land cover classes, we used an unsupervised learning approach to identify clusters of sites with similar land cover. We used k-means clustering (optimal *k* = 2 clusters) to identify sites that clearly differentiated in Principal Component Analysis (PCA) space, where the first PCA axis captured Okinawa’s primary land use gradient and explained 81.2% of the variance among our sites (Supplementary Figure S1). The PCA loadings show that the two clusters identified represent a distinction between sites that are primarily forested and those that are either agricultural or urban (Figures 1b and S1), hereafter together referred to as ‘developed’ sites.

**Figure 1.**
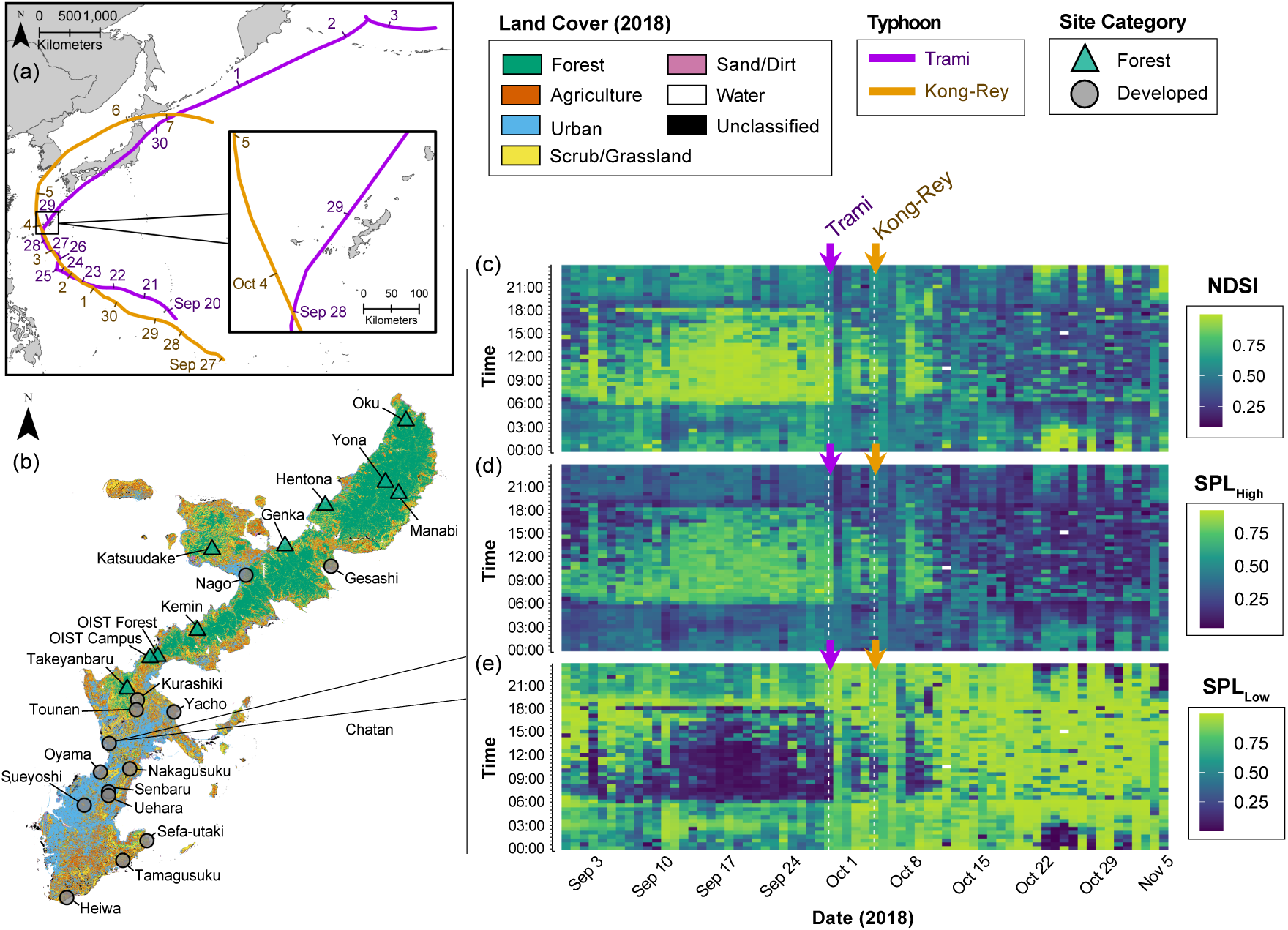
Field sites, timeline, and typhoon impact. (a) Map showing the tracks (coloured lines) of two large typhoons that hit Okinawa: super typhoon Trami in orange (20 Sep-03 Oct 2018; closest pass on 29 Sep 2018) and extratropical cyclone Kong-Rey in purple (27 Sep-07 Oct 2018; closest pass on 04 Oct 2018). (b) Map of Okinawa, including different land cover classifications based on a Landsat 8 image from 2015 (see Ross *et al*., 2018 for details). Twenty-four field sites with acoustic recorders are marked with coloured points; green triangles are sites grouped in the forested site cluster (*n* = 10), and grey circles those in the developed cluster (*n* = 14) based on unsupervised *k*-means clustering of land cover variables (see *Methods*). (c-e) Illustrative example time series of the study period at the developed Chatan field site (see b), showing the dates of the typhoon arrival (marked with coloured arrows) and the 30-day periods preceding and following typhoon impact. Dates along the X-axes span the full study period (30 Aug-04 Nov), and times along the Y-axes span 00:00-23:30 in half-hour intervals). Each grid cell then represents the value of a detrended and normalised acoustic index for the 10-minute recording corresponding to each time-by-date combination. To illustrate the potential for acoustic indices to reveal typhoon impacts, we show (c) the normalised difference soundscape index (NDSI), where higher values (lighter colours) represent a dominance of higher frequency sounds such as those attributable to biophony in the soundscape, while lower values (darker colours) comprise mostly low-frequency sounds such as anthropophony (Kasten et al., 2012), (d) sound pressure levels in the 2-11 kHz frequency range (SPL_High_), and (e) sound pressure levels in the 1-2 kHz frequency range (SPL_Low_). Sound pressure levels represent the amount of acoustic energy present in the higher (SPL_High_) or lower (SPL_Low_) frequency ranges and as such approximate the acoustic energy attributable to biophony and anthropophony, respectively (Kasten et al., 2012; see main text). Note the signal of the typhoons on the soundscape, clear in (d) as an increase in low-frequency sounds (*e.g.*, some geophony and anthropophony) relative to higher-frequency sounds following the typhoons. Geophony such as the heavy wind and rain caused by typhoons typically produces broadband sound, but the low-frequency range of the microphones saturated during the heavy geophony of the typhoons [see sharp increase in SPL_Low_ during the typhoon pass in (e)].

Acoustic data has been collected at each OKEON site since February 2017, but here we focus on a 66-day period in 2018 surrounding the landfall of two large typhoons, Trami and Kong-Rey. Trami passed closest to Okinawa on 29 September 2018 and was followed closely by Kong-Rey on 4 October 2018 (Japan Meteorological Agency [JMA] 2018; Figure 1a). We isolated recordings from the 30-day periods before (*pre-disturbance* period: 30 August – 28 September 2018) and after (*post-disturbance* period: 06 Oct – 04 Nov 2018) the typhoons made landfall, comprising a total of 771,840 minutes of data (Figure 1). Okinawa and the Ryukyu archipelago are increasingly exposed to more frequent and intense typhoons (A. Iwasaki, unpublished data), with annual typhoon seasons bringing disturbance events of varying magnitude (Elliott & Nino, 1960). Typhoon Trami (台風24号 in Japanese) was the largest typhoon (tropical cyclone) to hit Okinawa since OKEON acoustic recording began; it was Category 5-equivalent, with windspeeds reaching 183 km h^−1^ on 29 September 2018 (JMA 2018). Trami was followed shortly after by Kong-Rey (台風25号), which was less severe, striking Okinawa as a Category 3 extratropical cyclone (JMA 2018). Prior to these typhoons, the most recent large typhoon to pass Okinawa was Typhoon Neoguri (台風8号) in July 2014, so terrestrial habitats had likely recovered sufficiently to their climax communities for us to consider this an independent disturbance event. The chosen acoustic recordings therefore include a well-characterised pre-disturbance state (Figure S2; Ross *et al*., 2018, 2021a), followed by an extreme weather event and post-disturbance period during which soundscapes could potentially recover to their pre-disturbance state (Figures 1c and 1d). Though we did not test explicitly for effects of water-logging on microphone sensitivity following each typhoon, negligible effects of possible water-logging on microphone sensitivity are suggested by the fact that we find similar species vocalisation detection rates before and after the storm, and that SPL_High_ increases post-typhoon disturbance (see Results).

Okinawa’s soundscapes reflect a rich avifauna (McWhirter et al., 1996) including endemic, endangered, and habitat-restricted bird species (Inoue et al., 2019; Itô et al., 2000). As well as birds, anuran choruses are common at night, particularly during winter breeding seasons, though nighttime soundscapes tend to be dominated by orthopterans during the study period. Some cicadas are audible at this time of year, mainly during late morning after the onset of the dawn chorus, but not to the same intensity as during the peak summer months. Such biodiversity is generally higher in Okinawa’s forested north, while the southern field sites are more exposed to anthropophony (though most field sites are within audible distance of roads; Fig. 1; Ross et al., 2018). Many field sites are exposed to commercial and military air traffic, as well as fairly regular rainfall during the autumn months.

### Acoustic monitoring and data processing

Song Meter SM4 recorders (Wildlife Acoustics Inc., Concord, MA, USA) were installed at approximately breast height (~1.3m) at each field site and were programmed to record at default gain settings (+16 dB) via two omnidirectional microphones on a schedule of 10-minutes recording, 20-minutes standby, with recording starting on every hour and half hour. Data were saved to an SD card in stereo .WAV format at a sampling rate of 48-kHz. All audio data collected as part of the OKEON Churamori Project were archived with the Okinawa Institute of Science and Technology’s high-performance computing centre.

For each 10-minute audio file, we computed three commonly used acoustic indices in R (version 4.2.1; R Core Team 2022) using the *soundecology* package (version 1.3.3; Villanueva-Rivera & Pijanowski 2018). We calculated the normalised difference soundscape index (NDSI) by first generating a spectrogram via fast Fourier transformation (Hanning window size = 256) and splitting it into 1-kHz frequency bands. Sound pressure levels (SPL) were then calculated as the sum of the amplitude of all 1-kHz bands in the 2-11 kHz (SPL_High_) and 1-2 kHz (SPL_Low_) frequency ranges (Kasten et al., 2012) since these parameters were appropriate for our study system and objectives. The 2-11 kHz frequency range (summarised by SPL_High_) is typically dominated by biophonic sound sources in our system (including insects, birds, anurans, and other vocalising terrestrial animals), while the 1-2 kHz frequency range captured by SPL_Low_ is typified by low-frequency anthropophony (Kasten et al., 2012). NDSI is calculated as the ratio between these two components, such that higher values indicate a larger proportion of biophony in the soundscape relative to anthropophony; NDSI scales −1 to +1, where −1 indicates complete dominance of anthropophony (low-frequency sound) whereas +1 indicates total biophony (Kasten et al., 2012). This approach is preferable in our case over the original suggestion to compare anthropophony with the highest amplitude frequency band from the biophony range (Kasten et al., 2012), since it provides less weight to anthropophony and a greater focus on biophony (S. Gage, pers. comm.), which is important when considering biotic responses to typhoons. To facilitate comparisons among indices across studies, we normalised values of each acoustic index before measuring stability, producing relative proportions by dividing SPL_High_ and SPL_Low_ each by their site-specific maximum (Bradfer-Lawrence et al., 2020). We calculated NDSI using untransformed SPL_High_ and SPL_Low_ values, then normalised NDSI as (NDSI + 1)/2, since it ranges −1 and +1 and so cannot be scaled by its maximum to normalise values (Fairbrass et al., 2017).

Choice of acoustic indices was determined by previous work in this system showing that our focal indices were largely insensitive to potentially confounding sonic conditions such as the extreme geophony expected during typhoons (Ross et al., 2021a). These acoustic indices were not generally related to species richness in a previous study in this system (Ross et al., 2021a), nor to bird species vocalisation detections here (see below). However, SPL_High_ was positively correlated with the vocalisation detections of *Horornis diphone* when measured as site-level characteristics using mean acoustic index and vocalisation detections across the pre-typhoon period per field site (Bradfer-Lawrence et al., 2020). Moreover, though univariate acoustic features did not relate to bird point counts in a recent study, soundscape change was found to consistently reflect changes in community composition (Sethi et al., 2023). Accordingly, with well-described Okinawan soundscapes (Ross et al., 2018), and indices that are insensitive to acoustic masking by geophony (Ross et al., 2021a) and that have the potential to reflect ecological change or elements of biodiversity beyond simple bird vocalisations or richness, we concluded that these indices were fit for the purpose of revealing the impact of typhoon disturbance on Okinawa’s terrestrial ecosystems.

We also used machine learning methods (see Ross et al., 2018) to identify and count detections of three key focal bird species from our recordings. We used Kaleidoscope Pro (version 5.3.0; Wildlife Acoustics Inc., Concord, MA, USA) to train software recognisers for the large-billed crow (*Corvus macrorhynchos*, ハシブトガラス in Japanese), the Japanese bush warbler (*Horornis diphone*, ウグイス), and the Ryukyu scops-owl (*Otus elegans*, リュウキュウコノハズク). We chose these species because of their reliable detectability in recordings across sites; they are generally abundant, frequently vocalising, and we therefore had ample training data from which we built accurate species detectors with few false negative detections (though note that due to data volume, we did not explicitly quantify false negative rates). Our focal species represent a range of life histories, habitat affinities, and vocal repertoires (Hamao, 2013; Inoue et al., 2019; Itô et al., 2000; McWhirter et al., 1996; Ross et al., 2018), including a small-ranged forest habitat specialist (*O. elegans*), and are therefore expected to vary in their sensitivity to typhoons and land cover (see Table S1a for hypothesised typhoon effects on bird species detections). Species detection algorithms often transfer poorly across sites as a result of site-specific differences in background sonic conditions (Ross et al., 2018, 2021a), but we developed reliable detectors (≤15% false positives on visual inspection) at 21 sites for *C. macrorhynchos*, 17 sites for *H. diphone*, and 7 of the 10 forest sites for *O. elegans* (Table S2). Kaleidoscope Pro uses a supervised clustering approach based on Hidden Markov Models to separate sound types. Local experts cross-checked automated clustering of sound sources and reclassified sound clusters where necessary to refine species recognisers. Owing to the volume of data used in this study, we did not calculate exact false positive rates for species detections. Instead, we used Kaleidoscope Pro’s ‘distance-from-cluster-centroid’ measure to estimate identity confidence; larger distance values represent detections that are less likely to be the target species (see Pérez-Granados & Schuchmann 2020). Filtering by distance-from-centroid then allows rapid removal of low certainty detections. We chose a conservative distance filter of 0.5, though our results were qualitatively similar under less conservative filters (Figure S3). For all analyses, half-hourly bird detections were aggregated to daily detections to lessen the impact of individual singing events during the day (see Table S3).

### Analysis of acoustic indices

Before measuring the stability of acoustic indices through time, we detrended the normalised acoustic index time series using a centred moving average with a three-day window size in the *R* package *forecast* (version 8.14; Hyndman & Khandakar, 2008). We chose a three-day moving average because increasing the temporal window size of the moving average function to five or seven days produced qualitatively similar results at the expense of time series length and dampened soundscape dynamics (Figure S4). We then measured four components of stability at each site for normalised and detrended acoustic time series: temporal stability, resistance, recovery time, and spatial variability (Table S3; Donohue et al., 2013). Temporal stability was calculated as 1 minus the coefficient of variation (that is, the standard deviation divided by the mean) of the 30-day pre-typhoon period and the 30-day post-typhoon period, separately. Resistance was the maximum absolute change between the mean pre-typhoon baseline state and the maximum point of deviation from that state within 48 hours of the second typhoon passing, since resistance is a measure of the immediate to short-term responsiveness of the system (Hillebrand et al., 2018). Recovery time was 1 minus the time (in hours) between the point of maximum deviation from baseline (from which resistance was measured) and the point at which values returned to the pre-typhoon baseline (mean ± 95% confidence interval) and stayed within them for 24 hours (White et al., 2020), though results were generally robust to alternative window sizes (Figure S5). Temporal sampling regimes can affect the likelihood that meaningful ecological patterns are returned from acoustic datasets (Bradfer-Lawrence et al., 2019; Metcalf et al., 2021). To further validate our use of the entire 24-hr cycle in capturing ecologically meaningful results, we showed that mean daily acoustic index values were consistently positively related to those isolated from the dawn chorus (Figure S6), and those from non-dawn temporal subsets (Figure S7), regardless of field site.

At each time point, we calculated spatial variability as the coefficient of variation of acoustic index values for all 24 field sites (Table S3). Doing this per time point produced time series of spatial variability; spatial variability can change through time as the soundscapes at different field sites become more similar (lower spatial variability) or less similar (higher spatial variability), depending on a range of abiotic and biotic factors. Higher values of spatial variability among sites following a disturbance may represent a greater diversity of potential responses through asynchronous biomass fluxes within or among species, providing spatial insurance through patch dynamics (Leibold et al., 2004; Loreau et al., 2003; Wang et al., 2021). Spatial insurance effects occur via spatial exchange of individuals from different local patches in heterogeneous landscapes (Loreau et al., 2003). Since higher spatial variability represents greater heterogeneity, this, in turn, facilitates functional compensation or asynchronous dynamics among individuals or species in different local habitat patches with different environmental conditions, resulting in higher aggregate temporal stability (Wang et al., 2021). To test for potential land cover effects on spatial variability, we also calculated spatial variability among only those sites characterised as either forested or developed (Figure S1); that is, we separately measured the coefficient of variation of acoustic index values at the 10 forest sites and the 14 developed sites. This is because land cover change is expected to homogenise landscapes, reducing the opportunity for spatial insurance effects through asynchronous population dynamics (Wang et al., 2021). To aid comparison, stability components were normalised by their maximum (0-1) and defined such that larger values represent greater stability (see Table S1b for hypothesised typhoon effects, and Table S3 for detailed explanation of stability components and their interpretation).

All analyses were done in *R* (version 4.3.0, R Core Team 2023), using the packages *brms* and *segmented* (Bürkner, 2017; Muggeo, 2008). We tested for interactive effects of typhoons and land use on mean acoustic index states and temporal stability of indices, and for land use effects on acoustic index resistance and recovery time. For these analyses, we fitted generalised linear mixed effects models, with field site included as a random effect, using Stan (Stan development team 2020), implemented via the *brm* function in *brms* (Bürkner, 2017). For all four response variables, the modelled fixed effects included land use category (forest or developed) and typhoon state (pre- or post-typhoon). Given their nature, resistance and recovery time were not modelled as a function of typhoon impact. Default Hamiltonian Monte Carlo was used for the MCMC algorithm and all priors were uninformative. As our response variables fell on the [0,1] scale, we used the Beta model family with logit link. Model comparisons were made with leave-one-out cross validation (LOOIC) implemented in *brms* calculated via Pareto-smoothed importance sampling (Vehtari et al., 2017). We chose models with lowest LOOIC as best performing models (excepting cases where ΔLOOIC < 4.0, where model selection favoured the model with fewer parameters), since lower LOOIC indicates higher predictive accuracy. Four independent MCMC chains were run, each with a warmup phase of 5,000 iterations and sampling phase of 45,000 iterations. We inspected trace plots and density plots visually for chain mixture and verified convergence using the Gelman-Rubin R̂ < 1.01 and effective sampling size statistics (Gelman & Hill, 2006). We also tested for spatial autocorrelation of model residuals using the Moran’s *I* test statistic for each fitted model (Gittleman & Kot, 1990). Moran’s *I* results were always non-significant (that is, we did not detect significant spatial autocorrelation in any models), so we report results of the nonspatial models. Results of these models are presented as 95% highest density intervals (credible intervals) of all chains’ posterior parameter draws after the burn-in period.

For models of spatial variability responses, we fitted break-point models of spatial variability using the R package *mcp* (Lindeløv 2020). Break-point models fit segmented relationships between predictor and response variables to determine whether the form of the relationship changes as a function of the predictor variable. In our case, we modelled spatial variability as a function of time, with increasing numbers of break points from zero to three breaks, allowing intercepts, but not slopes, to vary between time segments. Break points were based on weakly informative prior distributions centring around the passing of each typhoon. That is, for a single break point, prior distributions centred on the mid-point between typhoons (around 1 Oct 2018), for two break points, priors were centred on each typhoon separately (29 Sep and 6 Oct 2018, respectively), and for three break points, we added a third point following the passing of the second typhoon (around 12 Oct 2018). For each model, we ran eight independent MCMC chains with a warmup phase of 10,000 iterations. We detrended acoustic index spatial variability time series with a periodicity of one week (this periodicity removed a suitable amount of seasonality and noise from spatial variability time series) using loess decomposition in the R package *stats* (R Core Team, 2023). We then ran all models on the trend time series (that is, removing the noise and seasonal components), since this provided a suitable trade-off between data resolution and model convergence. We compared models with different numbers of break points using LOOIC, again selecting models with lowest LOOIC values as best performing models unless a model with fewer breakpoints had ΔLOOIC < 4.0. Visual inspection of the number, location, and agreement of breakpoints within best performing models then reveals whether typhoons induced shifts in spatial variability; best performing models with strong agreement of breakpoints near the typhoons suggest a higher likelihood that observed changes in model intercept are driven by typhoons, and nonoverlapping 95% credible intervals between intercepts provides quantitative evidence for a change in intercept.

### Analysis of automated species detections

Given the lower temporal resolution of daily summed time series of bird species detections (*i.e.*, one value per day rather than 48), we did not estimate resistance or recovery time for bird species detections; such coarse grain time series would not produce meaningful resistance or recovery values in a time series of this length (particularly when defining a recovery window size for bird detections to be classed as recovered). Rather, we focused our analyses on the temporal stability of bird detections for each species across the 30-day pre- and post-typhoon periods and the spatial variability of detections per day across all sites, and across sites falling into each land use category (forested versus developed). Note that the forest specialist *Otus elegans* was not detected in any developed sites (Table S2), so for this species there is no data subset to compare between land cover types. As automated species detections produced count data, we did not normalise raw values of bird species detections.

As for acoustic indices, we tested for interactions between land use and typhoon effects on the mean number of daily detections (mean state) and the temporal stability of daily detections. We compared species effects by fitting a three-way interaction between species identity, land use, and typhoon period (two levels: before versus after the typhoons). We specified *brms* models as described previously, but with lognormal error distributions, which outperformed other error structures based on LOOIC. To aid convergence, we additionally set weakly informative priors of *N*(0,2) for all predictor variables in both models, but otherwise opted for uninformative priors. We measured spatial variability as the coefficient of variation of daily species detections across all 24 sites (Table S3), and again separately for the 10 forest sites and the 14 developed sites. For spatial variability, we fitted break-point models of spatial variability with intercept-only breaks, and used LOOIC to compare between models with zero to three breaks. Weakly informative priors followed break-point models of acoustic indices (above), and visual inspection of the number, location, and agreement of breakpoints within best performing models revealed whether typhoons induced shifts in spatial variability. Spatial variability values of species detections were already daily means, so we did not further detrend time series.

## Results

### Acoustic index results

We found that NDSI was significantly lower at many sites after the typhoons (Figure 2a). This overall pattern seemed not to be driven by an underlying change in 2-11 kHz sound pressure levels (SPL_High_; Figure 2b), but rather by an increase in SPL_Low_ following the typhoons (Figure 2c). There was no land use or typhoon effect on the temporal stability, resistance, or recovery of NDSI, SPL_High_, or SPL_Low_. However, in some cases we found differences in acoustic index values following the typhoons (Table S4).

**Figure 2.**
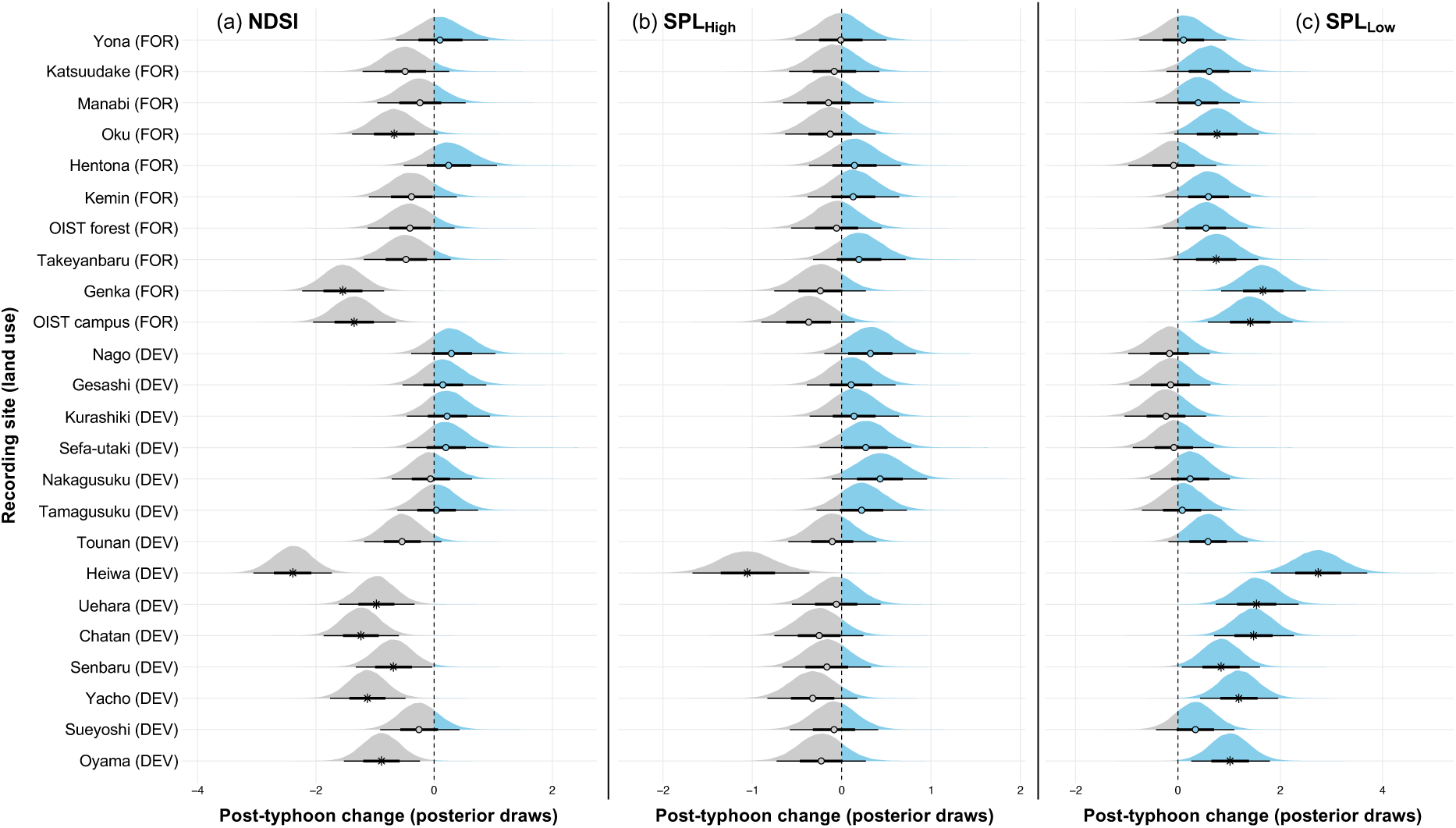
Comparison of acoustic index mean state values before and after the typhoons. Posterior distributions represent 90,000 post-convergence MCMC draws of the change from pre- to post-typhoon periods, where values below zero (grey) indicate a post-typhoon decline, and values above zero (blue) a post-typhoon increase in mean state value. Non-zero-spanning credible intervals are marked with *, while circles indicate zero-spanning credible intervals (no change based on the posterior distribution). Draws are shown per site, ordered from most forested (top) to most developed (bottom) based on principle component axis 1 of the land use dimensionality reduction (PCA; see Figure S1). Panels represent changes in mean state values for three normalised acoustic indices (see Methods): the normalised difference soundscape index [NDSI] (a), 2-11 kHz sound pressure levels [SPL_High_] (b), and 1-2 kHz sound pressure levels [SPL_Low_] (c).

When modelling the effects of typhoons on spatial variability of acoustic indices through time, models with three break points, including at least one coinciding with the typhoons, were chosen for all combinations of acoustic indices and land use (see Table S5). Following the typhoons, the spatial variability of NDSI increased (Figure S8). Break point models produced breaks that closely accompanied the typhoons, and this post-typhoon spatial divergence in NDSI was underlain by an increase in the spatial variability of sound pressure levels in the 2-11 kHz range (SPL_High_), but not in the 1-2 kHz range (SPL_Low_; Figures 3 and S9). Moreover, spatial variability in SPL_High_ was higher in developed than forested sites before the typhoons (compare “pre-typhoon” credible intervals in Table S5), and spatial variability among both groups of sites increased after the typhoons (Table S5; Figures 3b and S9). Conversely, the spatial variability of SPL_Low_ declined following the typhoons (Figures 3c and S9), but was not different between forest and developed sites (Table S5; Figure 3d).

**Figure 3.**
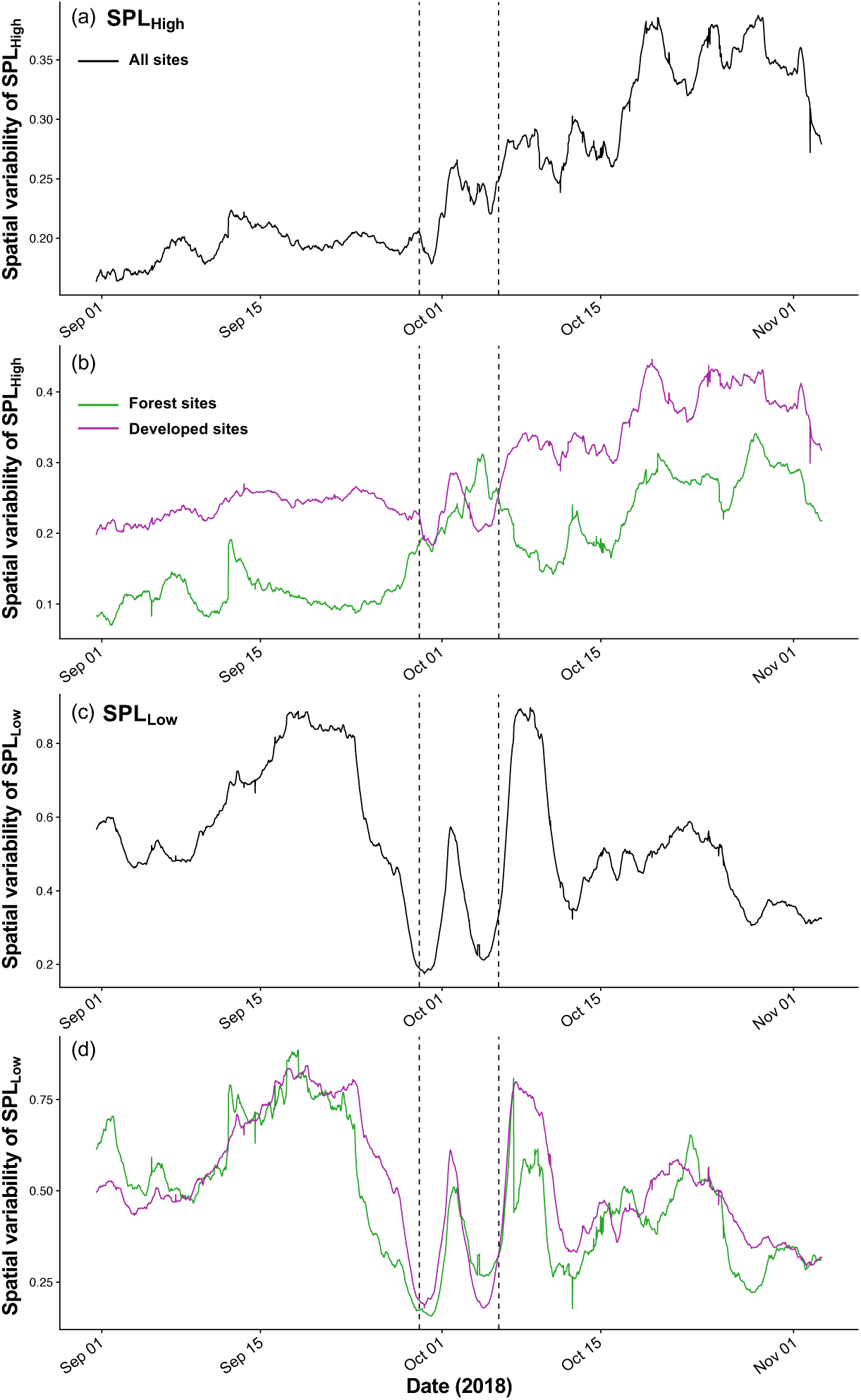
Spatial variability of 2-11 kHz [SPL_High_] and 1-2 kHz [SPL_Low_] sound pressure levels through time. Time series of SPL_High_ (a,b) and SPL_Low_ (c,d) spatial variability across all sites (a,c), and across forest (green) and developed (purple) sites separately (b,d). Dashed lines delineate the pre- and post-typhoon periods.

### Automated species detection results

Species identity interacted with the typhoons, producing species-specific typhoon responses (Table S4; Figure S10). Detections of *C. macrorhynchos* and *O. elegans* were similar preceding and following the typhoons (Figures 4a and 4c), whereas *H. diphone* was detected less often after the typhoons (Figures 4b and S11). We also found that, following the typhoons, species detections were more stable (less variable) through time, regardless of the species considered (Figure 5; Table S4). We found no effect of land use on the mean number of daily species detections or the temporal stability of daily detections (Table S4; see Figure S10 for a map of mean daily detections before and after the typhoons).

**Figure 4.**
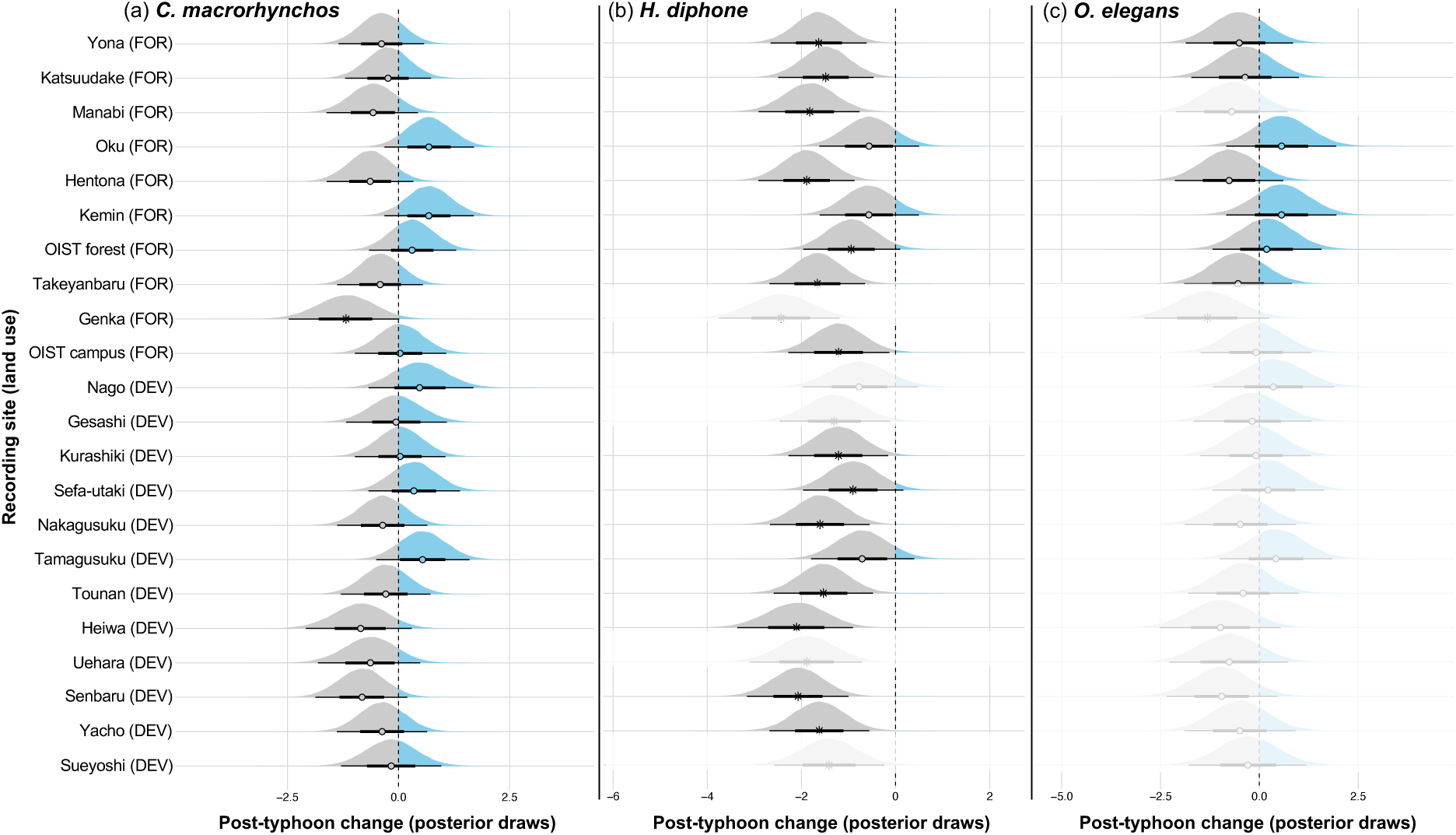
Comparison of mean daily species detections before and after the typhoons. Posterior distributions representing 90,000 post-convergence MCMC draws of the change from pre- to post-typhoon periods, where values below zero (grey) indicate a post-typhoon decline, and values above zero (blue) a post-typhoon increase in mean daily species vocalisation detections. Non-zero-spanning credible intervals are marked with *, while circles indicate zero-spanning credible intervals (no change based on the posterior distribution). Draws are shown per site, ordered from most forested (top) to most developed (bottom) based on principle component axis 1 of the land use dimensionality reduction (PCA; see Figure S1). Panels represent changes in mean daily species detections for our three focal species: *Corvus macrorhynchos* (a), *Horornis diphone* (b), and *Otus elegans* (c). Inferred posterior draws (automatically computed through the site random effect term) extrapolated to field sites where species were not present (Table S2) are shown as faded distributions.

**Figure 5.**
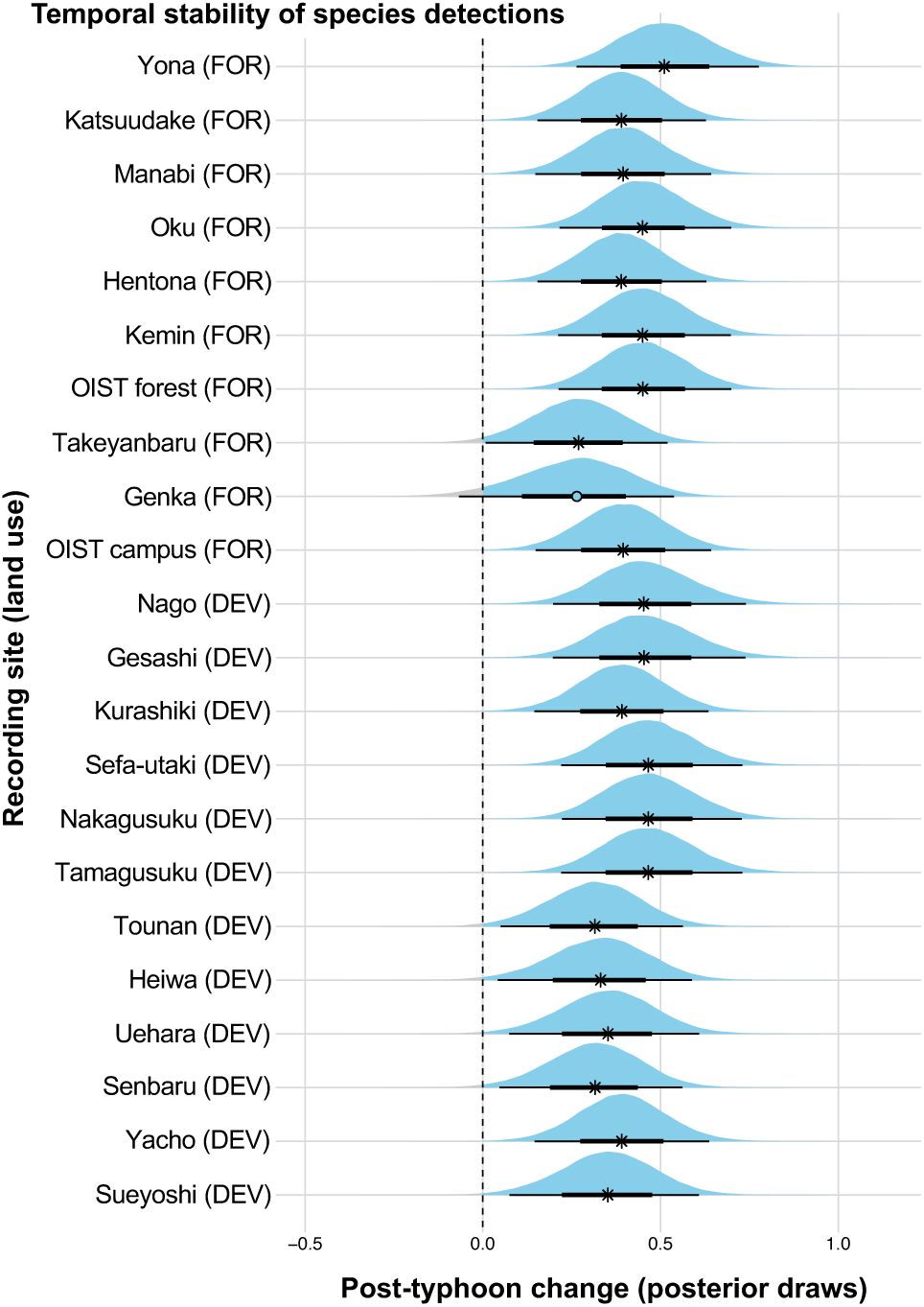
Comparison of temporal stability of species detections before and after the typhoons. Posterior distributions represent 90,000 post-convergence MCMC draws of the change from pre- to post-typhoon periods, where values below zero (grey) indicate a post-typhoon decline, and values above zero (blue) a post-typhoon increase in the temporal stability of automated species vocalisation detections (all species). Non-zero-spanning credible intervals are marked with *, while circles indicate zero-spanning credible intervals (no change based on the posterior distribution). Draws are shown per site, ordered from most forested (top) to most developed (bottom) based on principle component axis 1 of the land use dimensionality reduction (Figure S1). See Figure S12 for posterior draws of individual species.

When modelling the effects of typhoons on spatial variability of bird detections through time, we found species-specific typhoon effects, with detections of *Corvus macrorhynchos* and *Otus elegans* not showing clear typhoon-related changes in spatial variability (Table S5; Figures 6a and S13). However, across forested sites, detections of *C. macrorhynchos* showed an increase in spatial variability coincident with the typhoons (as indicated by the selected break-point model, Figure S13b). The spatial variability of *Otus elegans* detections was higher than of the other bird species (Table S5; Figure 6a). In contrast to the other species, detections of *Horornis diphone* became more variable around the onset of the typhoons across all sites (Figure 6a) and across forested sites (Figure 6b), as supported by break-point models (Figure S13d and S13e). However, across developed sites, there was no clear association between the typhoons and any possible break points in *H. diphone* spatial variability (Table S5; Figures 6b and S13f).

**Figure 6.**
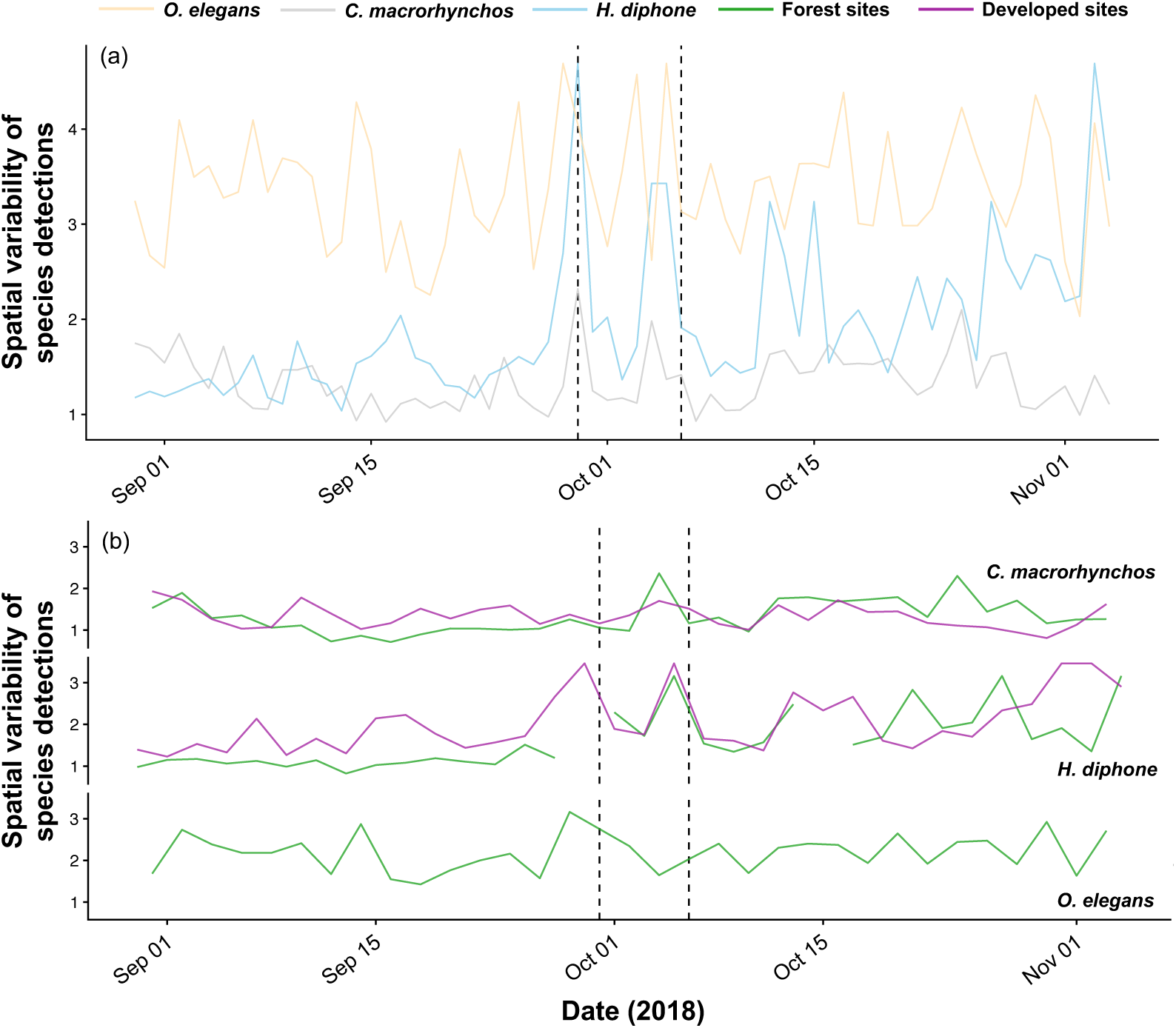
Spatial variability of automated daily species detections through time. Time series of spatial variability (coefficient of variation) of daily species detections across all sites (a) for each of *Corvus macrorhynchos* (grey), *Horornis diphone* (blue), and *Otus elegans* (yellow), and (b) measured separately across the 10 forest (green) and 14 developed (purple) sites for each species. Dashed lines delineate the pre- and post-typhoon periods.

## Discussion

This study leverages high-resolution acoustic monitoring data from an island-wide sensor array to record ecological responses to extreme weather events in the form of two large typhoons. We found no land use effects on most dimensions of stability measured. However, we found that spatial variability of the 2-11 kHz sound pressure levels (SPL_High_, a proxy for biophony; Kasten et al. 2012) was greater among developed sites compared with forested ones before the typhoons, and that the spatial variability of both SPL_High_ and the normalised difference soundscape index (NDSI) increased significantly following the typhoons. Spatial variability continued to increase at the end of the study, which may be reflective of ecological responses to typhoons that extend beyond the immediate and short-term responses we aimed to document in this study. Contrary to the expected typhoon-induced spatial homogenisation of soundscapes, our results indicate that the typhoons elicited divergent ecological responses among Okinawa’s soniferous animal communities. In support of this, we found that detections of two of our three focal bird species, *Corvus macrorhynchos* and *Horornis diphone*, were more spatially variable after the typhoons (though *C. macrorhynchos* detection spatial variability increased only among the forested sites), in contrast to the expected post-typhoon homogenisation through mortality or distributional shifts. This post-typhoon increase in spatial variability of species detections was present among forested but not developed sites, suggesting perhaps that land use and habitat change can hinder the reactive capacity of ecological communities and their associated soundscapes. That our species-level spatial variability results matched those of acoustic indices further validates our use of acoustic indices to assess typhoon impacts in Okinawa.

We found spatial divergence of NDSI and SPL_High_, but not SPL_Low_ or species detections, in developed urban and agricultural sites. This disconnect between results from the SPL_High_ index versus automated species detections in developed sites could reflect differences in local biodiversity in developed sites, or perhaps that differences in ambient sonic conditions affected our ability to accurately detect species responses (Ross et al., 2021a). We did not directly measure local biodiversity in this study, instead estimating the activity of some key focal bird species using automated species detections. However, previous work in this system provided evidence for a loss of rare and endemic birds under land use development, producing communities that are a nested subset of forest bird communities in Okinawa’s developed south (Ross et al., 2018). The species surveyed here did not exhibit responses that diverged in space in developed sites following the typhoons, as might have been expected based on SPL_High_ results in developed sites. This suggests that the spatial divergence in the biotic component of soundscapes recorded here may be better explained by other species (of birds or other taxa) not targeted in this study. Future work expanding on these analyses to provide a more holistic view of the Okinawan biota should therefore prove fruitful for identifying individual species contributions to typhoon responses. If our SPL_High_ results are indeed a product of biotic responses to typhoons as would be expected from theory (Kasten et al., 2012), and as supported by our finding that spatial variability of both *C. macrorhynchos* and *H. diphone* increased among forested sites, then a post-typhoon increase in spatial variability may reflect changes to species’ patchiness. For example, Willig and Camilo (1991) described an increase in spatial patchiness of the snail *Caracolus caracol* following Hurricane Hugo in Puerto Rico, caused by a thinning of populations due to post-hurricane mortality.

Soundscape composition after the typhoons saw an increase in low-frequency sound pressure levels (SPL_Low_) as anticipated (Table S1b), but not a decline in SPL_High_ as might be expected were animal populations impacted negatively by typhoon disturbance (*e.g*., Cely, 1991; Pavelka et al., 2007). In contrast, the observed post-typhoon increase in spatial variability in NDSI was driven by SPL_High_ rather than SPL_Low_. This suggests that, while SPL_High_ may not have been affected substantially by typhoon disturbance at the individual site level (Figure 2b), variation in biotic responses at larger scales across field sites nonetheless manifested as changes in the spatial variability of SPL_High_ after the typhoons. This is likely driven, at least in part, by the observed post-typhoon spatial divergences in species vocalisations. That we did not detect particularly strong site-level typhoon impacts on soundscapes, but rather saw spatial divergence in ecological responses to typhoons across multiple sites, underscores the necessity and benefits of monitoring at scale. Multi-site acoustic sensor arrays such as ours thus provide opportunity to monitor both local and regional biodiversity change, in turn providing critical new insight for conservation management (Roe et al., 2021; Sethi et al., 2020a; Van Parijs et al., 2015). The observed post-typhoon increase in SPL_Low_ on the other hand, likely reflects increased human activity at developed sites as efforts to survey and repair typhoon damage got underway (S. Ross pers. obs.). Conversely, the expected post-typhoon increase in SPL_Low_ observed at a few forest sites may reflect changes in sound propagation driven by vegetative structural damage and thinning of previously dense habitats, as is often documented following large storms (Abbas et al., 2020; Elliott & Nino, 1960). We did not measure habitat structure directly, and so the causes of increases to SPL_Low_ following typhoons Trami and Kong-Rey cannot be demonstrated empirically. We did, however, observe significant damage and alterations to habitat structure at our forested field sites (T. Yoshida & M. Yoshimura, pers. obs.). Automated bird species detections were more stable through time after the typhoons, suggesting disturbance may affect the consistency of species vocalisations in Okinawa (see also Fraterrigo & Rusak, 2008), as might be expected if rare or infrequent vocalisers are lost from soundscapes (Bradfer-Lawrence et al., 2020; Table S1b), or perhaps that typhoon-induced changes to habitat structure allow vocalisations to travel further without attenuation, and hence be detected more reliably by our sensors. That automated bird species detections were more stable through time after the typhoons but the temporal stability of sound pressure levels in the higher frequency range (approximating biophony; Kasten et al., 2012) did not change after the typhoons suggests that these SPL_High_ measures reflect other aspects of the biotic soundscape beyond those three target species included in our analysis.

The focal bird species considered here differed in their responses to typhoons. Automated vocalisation detections of the Japanese bush warbler (*Horornis diphone*) declined after the typhoons, while, as expected, those of the large-billed crow (*Corvus macrorhynchos*) did not. Contrary to expectation (Table S1a), however, detections of the Ryukyu scops owl (*Otus elegans*) were also unaffected. Given that acoustic surveys cannot differentiate between cases where a species is not producing sound and those where that species is not present (Toth et al., 2022), we cannot say with certainty that *H. diphone* populations declined following the typhoons. Regardless, our detected post-typhoon declines in *H. diphone* vocalisations—either through behavioural changes, distributional shifts, or local mortality—were consistent across >80% of the field sites in which this species was detected. We expected that insectivores, such as *H. diphone*, may not be adversely affected by the typhoons (excepting effects of direct mortality), but rather that structural changes to habitat may increase prey availability in canopy gaps (Cely, 1991; Seki, 2005). However, for *H. diphone* we observed the opposite pattern. Azuma *et al*. (1997) documented considerable effects of undergrowth removal on invertebrate communities in Okinawa. Given that *H. diphone* relies on undergrowth and bushes for foraging (Haneda & Okabe, 1970) and typhoon disturbance has the potential to alter the structure of this habitat (Abbas et al., 2020; Elliott & Nino, 1960), the observed decline in *H. diphone* detections following the typhoon may therefore be due to bottom-up changes to the invertebrate communities on which *H. diphone* feeds following typhoon damage to the undergrowth (Azuma et al., 1997). Moreover, *H. diphone* is a small-bodied species and may thus be especially vulnerable to weather-related mortality (Brown & Brown, 1998). In contrast, and contrary to expectation (Table S1a), the forest specialist *O. elegans* was not detected less frequently after the typhoons, suggesting that its habitat or foraging were unaffected by the typhoons, or perhaps that cavity nesting reduced typhoon impact by reducing exposure to the extreme weather (Inoue et al., 2019). Overall, species detections were more stable through time after the typhoons, though acoustic index temporal stability was not affected by typhoons. Given that soundscapes can be more temporally stable when human activity is restricted (Ross, 2022), the higher post-typhoon temporal stability of species detections observed here may relate to altered human space use across Okinawa following the typhoons.

Species-specific responses to disturbance may more generally reflect differences in life history and other functional response traits (Suding et al., 2008), which can be useful predictors of community dynamics, disassembly, and stability in birds (*e.g.*, Ausprey et al., 2022; Hordley et al., 2021; White et al., 2023). Similarly, different typhoon responses of vocalisation detections among field sites may reflect differences in underlying vegetative changes as determined by plant functional response traits. For example, Craven *et al*. (2016) found that functionally diverse Canadian forests were dominated by trees with response traits that promoted resilience to recurrent anthropogenic disturbance through rapid regrowth, rather than resistance to projected climate change through drought or flood tolerance. The response traits of plants may then, in turn, determine the structural habitat change experienced by birds and other vocalising animals (*e.g.*, Abbas et al., 2020), as well as directly influencing sound propagation (Morton, 1975).

Though we and others have demonstrated the capacity for passive acoustic monitoring methods to capture unpredictable extreme weather events (Gottesman et al., 2021; Simmons et al., 2021), such methods are often limited in their ability to accurately reflect biodiversity patterns. A recent meta-analysis reported a generally positive link between acoustic indices and biodiversity (Alcocer et al., 2022), but one with diminishing effect sizes over time, as studies increasingly forego appropriate validation and as their designs incorporate yet wider varieties of non-target sounds, which can hinder the interpretability of those acoustic indices aiming to reflect biodiversity (Ross et al., 2021a). Moreover, Sethi *et al*. (2023) recently reported that univariate acoustic features (akin to acoustic indices) did not well reflect bird species richness across four datasets from India, Malaysia, Taiwan, and the United States. However, soundscape change was nonetheless found to consistently reflect changes to underlying bird community composition, suggesting soundscape dynamics are useful proxies for ecological dynamics. The authors conclude by suggesting acoustic features are used in combination with traditional field surveys or other methods of ground-truthing soundscape data (Sethi et al., 2023). Similarly, Müller *et al*. (2023) used acoustic indices and deep learning methods to monitor biodiversity recovery after agricultural abandonment in an Ecuadorian rainforest. Though acoustic indices may not well reflect biodiversity, they nevertheless reflect biodiversity dynamics in a range of systems.

The joint use of acoustic indices and automated species detections in our study provides two separate lines of evidence for typhoon-induced soundscape change. Such species and soundscape methods are still rarely used in combination despite their clear potential to provide complementary information on ecological dynamics (*e.g.*, Ferreira et al., 2018; Ross et al., 2018). Indeed, our complementary results linking post-typhoon spatial divergence of SPL_High_ to that of species detections underscore the efficacy of combing both acoustic indices and individual species recognition models. That said, building reliable vocalisation recognition algorithms remains a challenge, particularly when aiming for transferability to different habitats or seasons, which provide a range of non-target sounds beyond those on which algorithms may have been trained. Increasing application of deep learning to such problems will likely help provide a solution (*e.g.*, Sethi et al., 2020b) as will continued efforts to build labelled sound libraries from which automated species detection algorithms can be trained (Deichmann et al., 2018).

Soundscape dynamics are frequently characterised by strong seasonal cycles (*e.g.*, Vokurková et al., 2018). This presents a challenge when attempting to disentangle disturbance responses from seasonal soundscape change. For example, our focal species differ in their seasonality and phenology, meaning that natural phenological differences may be in part responsible for the differences in species’ typhoon responses we observed here. A recent study of Okinawan ants showed that ant activity was more seasonal in forested than developed field sites, and that these patterns in seasonality were driven by asynchronous seasonal dynamics of different ant species at the community level (Kass et al., 2023). This raises the possibility that land development degraded seasonal dynamics in the soundscape, possibly contributing to differences in the spatial divergence of SPL_High_ among forested but less so among developed field sites. However, in our study, *Horornis diphone* vocalisation detections declined sharply and concomitantly with the typhoons (Figure S11). This suggests the observed post-typhoon decline was most likely a result of the typhoons themselves, rather than a seasonal decline, which we would expect to be much more gradual in nature. Our moving average detrend aimed to remove as much seasonal signal as possible, though more sophisticated approaches to deseasonalisation exist such as, for example, wavelet decomposition. Such approaches typically require longer time series than ours, however, with clear periodic components comprising less than a quarter of the total time series length (Cazelles et al., 2008), and so wavelet analysis was not appropriate for our study. As the OKEON Churamori Project continues to collect soundscape recordings from across Okinawa, advanced time series models and decomposition techniques will be needed to process yet longer time series and periodic signals.

Our *k*-means clustering approach to distinguish field sites by their dominant land use identified an optimal split of two clusters, separating primarily forested sites from those dominated by developed urban or agricultural land use. However, these developed land uses can act on ecological dynamics and stability in different ways. For example, Olivier *et al*. (2020) used citizen science data from across France to show that agricultural intensification directly affected population, and, in turn, community stability of birds, whereas urbanisation acted only indirectly on community stability through changes to diversity and population asynchrony. Our study design, which was based on unsupervised (*k*-means) site clustering by dominant land use, consequently did not allow us to directly compare urban and agricultural field sites, despite their potential for contrasting effects on ecological stability. Moreover, we classified sites based on land cover from a 2015 Landsat 8 image, while habitat structure and land use are dynamic features. As such, we could not directly capture legacy effects of historic land use on ecological communities in this study, including both intensification—the dominant trend in Okinawa—and abandonment, which can each act as driving forces structuring biodiversity (Daskalova et al., 2020; Daskalova & Kamp 2023).

## Conclusion

Our study tested the capacity for land use and climate change in the form of extreme weather events to jointly shape ecological stability. Using passive acoustic monitoring data from a landscape-scale sensor network across Okinawa Island, we found that land use rarely modified ecological responses to typhoons. However, soundscapes diverged across the landscape following the typhoons, contrary to the expected typhoon-induced soundscape homogenisation. The 2-11 kHz sound pressure levels (SPL_High_) in developed sites was initially more variable across sites, but this post-typhoon spatial divergence occurred among both forested and developed urban and agricultural field sites. However, species detections diverged in space among forested but not developed sites following the typhoons, suggesting a wider variety of potential disturbance responses among forest bird assemblages than those in developed sites. That is, land use intensification may produce ecological communities that are more homogeneous in how they respond to disturbance (Vogel et al., 2019), while forest sites harbour communities with greater potential for collective resilience to future disturbance through patch dynamics and rescue effects among different local forest communities (Leibold et al., 2004). Such spatial insurance effects have the potential to contribute to landscape-scale stability and spatial portfolio effects by affecting population and community asynchrony (Loreau et al., 2003; Wang et al., 2021). Further studies of disturbance responses in Okinawa should aim to explicitly identify such spatial insurance effects for birds or other taxa.

This study draws on prior knowledge of Okinawan biodiversity (Inoue et al., 2019; Itô et al., 2000; McWhirter et al., 1996), the performance of passive acoustic methods in this system (Ross et al., 2018, 2021a), and the characteristics of typhoons and land use intensification across Okinawa Island (Elliott & Nino, 1960; Takeuchi et al., 1981), to explore fundamental ideas concerning disturbance and stability (Ross et al., 2023). Such baseline data provides a critical backdrop against which our results stand, allowing us to infer species and soundscape responses to the joint threats of climate change and land use intensification from acoustic recordings of typhoons (Altwegg et al., 2017). As longer and higher-resolution acoustic data are amassed through multi-site acoustic sensor arrays (*e.g.*, Roe et al., 2021; Sethi et al., 2020a; Van Parijs et al., 2015), the utility of passive acoustic monitoring to document ecological responses to extreme weather events across the globe will become ever clearer, particularly in light of the increasing frequency and destructive potential of extreme events in the Anthropocene.

## Acknowledgements

Many individuals in the OKEON Churamori Project contributed to data collection, site maintenance, and community outreach; we especially thank Masako Ogasawara, Mayuko Suwabe, Shinji Iriyama, Toshihiro Kinjo, Izumi Maehira, Yuko Matsudo, Seiichiro Nakagawa, Shoko Suzuki, Takumi Uchima, Madoka Oguro, Dan Warren, and Chisa Oshiro. We are grateful for the help and support provided by the Scientific Computing and Data Analysis section of the Research Support Division at OIST. Our sincere thanks go to the landowners, museums, local governments, and schools that host the OKEON Churamori Project field sampling sites, and to the people of Okinawa. We also thank Marina Lašić and Rob McHenry for their contributions to building automated species recognisers, Yvonne Buckley, Anne Magurran, and Jamie Kass for helpful discussion, and Vito Muggeo for advice on break-point modelling in R. This work was supported by subsidy funding to OIST, and an Irish Research Council Postgraduate Scholarship [GOIPG/2018/3023] and Canon Foundation in Europe 2021 Research Fellowship awarded to S.R.P-J.R.

## Conflict of Interest

The authors declare no conflict of interest.

## Data availability statement

The data supporting the findings of this study and all R code are available via the Zenodo digital repository as Ross, S. R. P.-J., Friedman, N. R., Dudley, K. L., Yoshimura, M., Yoshida, T., Economo, E. P., Armitage, D. W., & Donohue, I. (2023). Data from: Divergent ecological responses to typhoon disturbance revealed via landscape-scale acoustic monitoring (v0.3-review2). *Zenodo* https://doi.org/10.5281/zenodo.8133339

## CRediT Author Contribution Statement

SRP-JR: Conceptualisation, Methodology, Software, Validation, Formal analysis, Investigation, Writing – Original Draft, Writing – Reviewing & Editing, Visualisation, Funding Acquisition; NRF: Conceptualisation, Software, Resources, Data Curation, Writing – Reviewing & Editing, Project Administration; KLD: Software, Validation, Resources, Data Curation, Visualisation, Project Administration; TY: Resources, Data Curation, Project Administration; MY: Resources, Data Curation, Project Administration; EPE: Resources, Data Curation, Writing – Reviewing & Editing, Project Administration, Funding Acquisition; DWA: Methodology, Software, Formal Analysis, Visualisation, Writing – Reviewing & Editing; ID: Conceptualisation, Methodology, Writing – Original Draft.

## Notes

https://doi.org/10.5281/zenodo.8133339

### Competing Interest Statement

The authors have declared no competing interest.

### Summary of Updates

Interpretation of sound pressure levels has been toned down, plus minor text edits throughout, and revisions to figures for clarity. Additional text in discussion surrounding correspondence between bird detection results and acoustic index results, as well as what acoustic index values reflect biologically.

