## Supplementary Information for "Divergent ecological responses to typhoon disturbance revealed via landscape-scale acoustic monitoring"

**Table S1. Expected typhoon impacts. (a)** Life histories and characteristics of our three focal bird species, and hypothesised effects of typhoons on species based on their characteristics. **(b)** Hypothesised effects of typhoons on the different variables included in this study. Note NDSI results may be driven by $SPL_{High}$ and/or $SPL_{Low}$, depending on the strength of the effect for each component index.

| (a) Focal bird species | *Diet (foraging)* | *Habitat affinity* | *Vocalisation properties* | *Expected typhoon impact* |
|---|---|---|---|---|
| *Corvus macrorhynchos* | Generalist (generalist). | Generalist, urban. | Loud “Caw” ~1 – 3 kHz with harmonics up to ~8 kHz. | *Minimal impact*. Generalist diet and habitat affinity mean *C. macrorhynchos* should be flexible and able to adapt to changes in diet or habitat following typhoon disturbance. |
| *Horornis diphone* | Small invertebrates and larvae (undergrowth). | Dense bush / undergrowth. | Long single note (~1.5 kHz) with trill up to ~4 kHz. | *Moderate – Severe impact*. Typhoon damage to undergrowth may impede nesting and foraging. Small body size likely makes *H. diphone* vulnerable to weather-related mortality (Brown & Brown 1998). |
| *Otus elegans* | Large arboreal invertebrates, especially orthopterans (generalist). | Forest specialist (old growth tree cavities). | Single or double “hoo” ~1 kHz. | *Moderate impact*. Typhoon damage to forest structure may reduce prey substrate and availability, or increase invertebrate prey access in canopy gaps (Cely 1991). |

| (b) Variable | Expected typhoon impact: | | |
|---|---|---|---|
| | ***2-11 kHz Sound Pressure Level*** *($SPL_{High}$)* | ***1-2 kHz Sound Pressure Level*** *($SPL_{Low}$)* | ***Bird detections*** |
| *Index or Detection Vlaues* | *Post-typhoon decline*. Typhoons should reduce biophony by direct mortality or relocation of vocalising animals (Wiley & Wunderle 1993). | *Post-typhoon increase*. Damage to canopy structure may produce forest gaps through which higher anthropophony can be heard (Cely 1991), this effect should be less in developed sites. | *Post-typhoon decline*. Typhoons should reduce species detections by direct mortality or relocation of vocalising animals (Wiley & Wunderle 1993), but species should differ in their responses (Table S1a). Greater potential structural change and sensitivity of specialist species in forests should cause larger declines in forest sites than developed sites. |
| *Temporal Stability* | *Post-typhoon increase*. Reduced density of vocalising animals should produce more consistent (stable) soundscapes, as rare or infrequent vocalisers are lost (Bradfer-Lawrence et al. 2020), particularly in forest sites. | *Minimal change*. Changes to forest structure following typhoons may increase anthropophony propagation, but anthropophony is anyway infrequent (low stability). There is no expected change in developed sites. | *Post-typhoon increase*. Reduced density of vocalising animals should produce more consistent (stable) soundscapes, as rare or infrequent vocalisers are lost (Bradfer-Lawrence et al. 2020), particularly in forest sites. |
| *Resistance / Recovery* | *High resistance, slow recovery*. Closed canopy forests should provide greater resistance than developed sites (Abbas et al. 2020), but greater potential structural habitat change and mortality should slow forest recovery (Elliott & Nino 1960). | *Low resistance, fast recovery*. Broadband geophony during typhoons may be recognised as anthropophony. Open habitat structure in developed sites should produce low resistance, but limited structural change should result in faster recovery compared to forest sites (Raymond et al. 2020). | - |
| *Spatial Variability* | *Post-typhoon decline*. Typhoons should homogenise soundscapes by removing specialists and damaging habitat structure, particularly in forest sites (Cely 1991). | *Post-typhoon decline*. Typhoons should homogenise soundscapes by damaging habitats and opening them to anthropophony in forest sites (Raymond et al. 2020). Lower expected structural damage in developed sites should mean less severe homogenisation than in forests sites. | *Post-typhoon decline*. Typhoons should homogenise species detections across forest sites due to change in habitat structure and direct mortality or distribution shifts (Wiley & Wunderle 1993). Species should be differently affected (Table S1a), and because developed sites should suffer less structural habitat change, they should be less homogenised. |

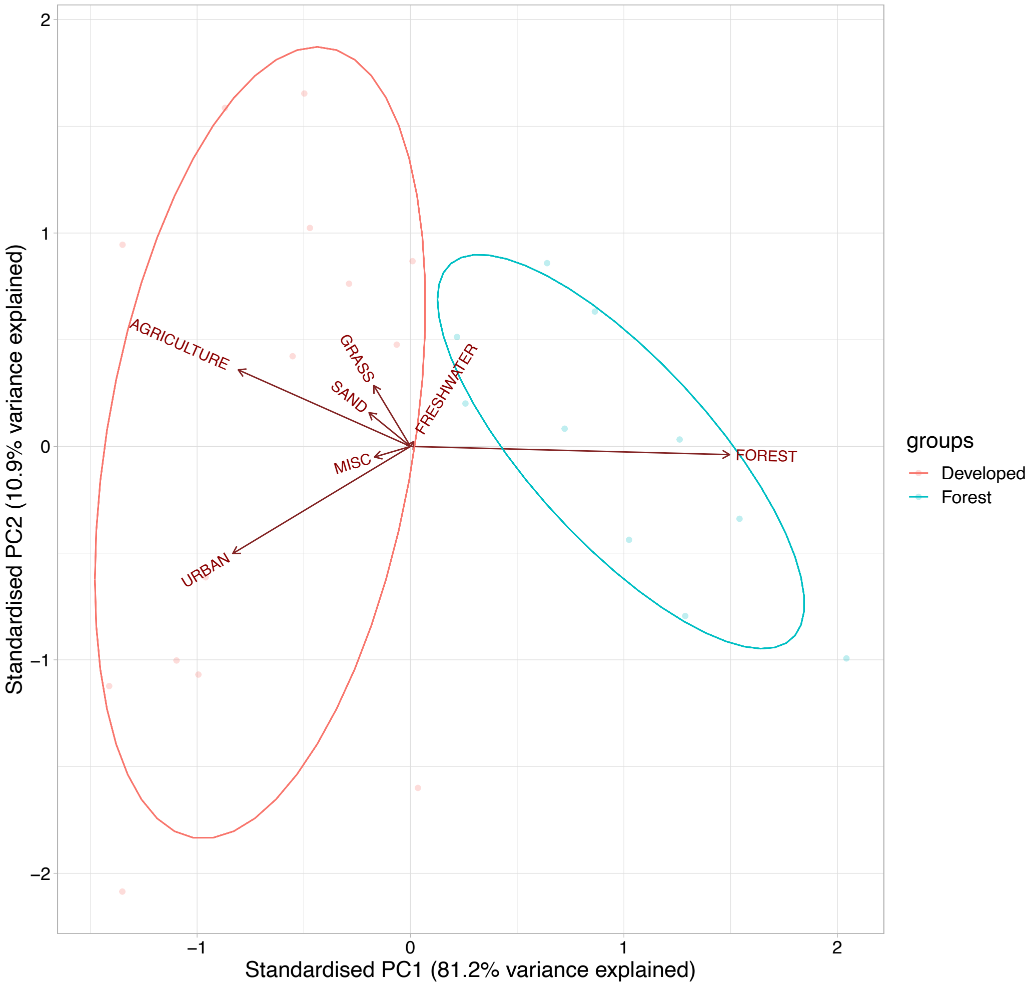


**Figure S1. Ordination biplot for the land cover variables used in this study**. We used unsupervised k-means clustering (optimal *k* value = 2 clusters) of a Principal Component Analysis (PCA) of land cover variables (see Methods) to automatically identify clusters of sites with similar land cover. We found a clear distinction between two clusters (red versus blue points and ellipsoids). When examining the variable loadings (land cover classes marked on PCA ordination with names and arrow length showing relative variable weights), we found that the clusters represented a clear distinction along PCA axis 1 (variance explained = 81.2%), where the 10 blue sites were primarily forest sites, while the 14 red sites with either primarily agricultural, urban, or managed grassland sites, herein collectively termed 'developed' sites.

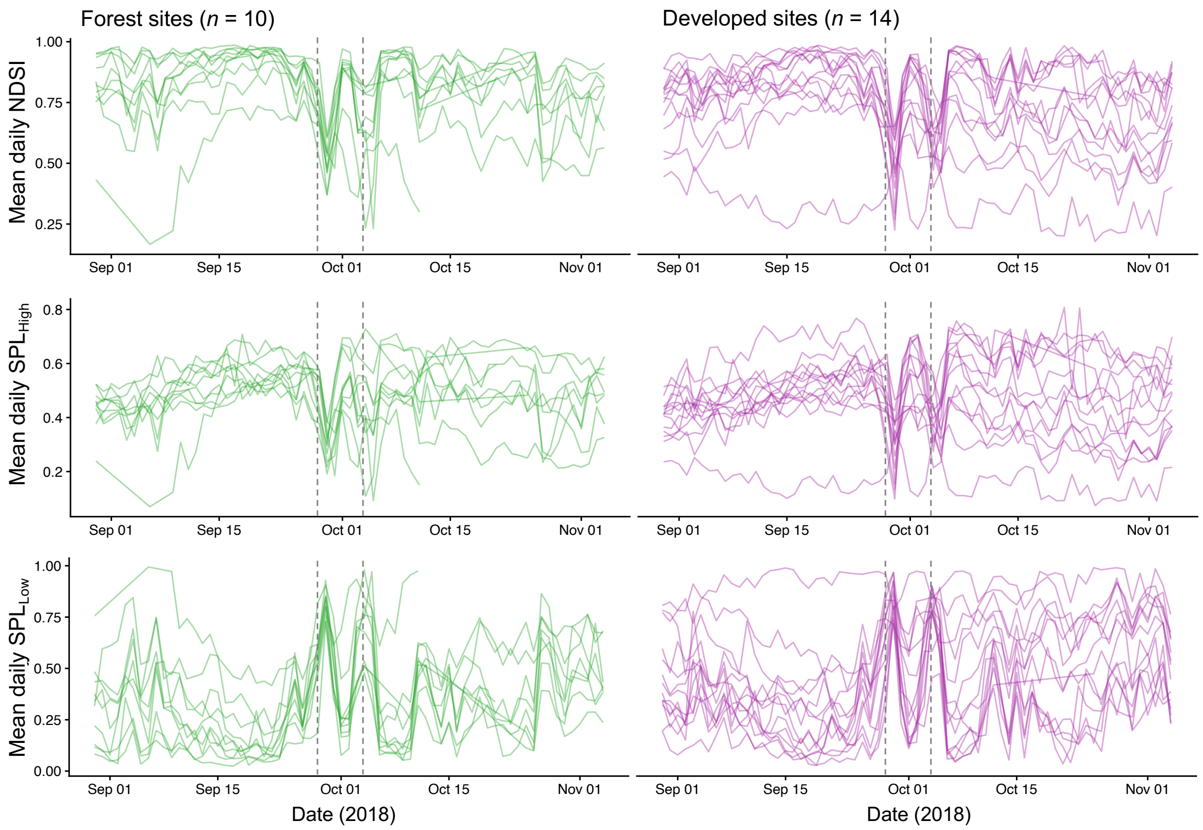


**Figure S2. Acoustic index time series**. Time series are aggregated to daily mean acoustic index values for visualisation purposes and separated by land use category based on *k*-means clustering (Figure S1), with the 10 forest field sites on the left (green) and 14 developed field sites on the right (purple). Y-axis values represent three acoustic indices: the Normalised Difference Soundscape Index (top), 2-11 kHz sound pressure levels [$SPL_{High}$] (middle), and 1-2 kHz sound pressure levels [$SPL_{Low}$] (bottom). Grey dashed lines represent the closest pass of the two large typhoons: super typhoon Trami (29 Sep 2018) and extratropical cyclone (04 Oct 2018).

**Table S2. Automatic supervised learning bird species detection classifiers across field sites**. We produced automated (supervised machine learning) species vocalisation classifiers in Kaleidoscope Pro (version 5.3.0; Wildlife Acoustics Inc., Concord, MA, USA), using training data from across a variety of sites and dates, and applying trained classifiers to the full dataset of recordings (30 Aug-04 Nov 2018) to automatically identify species detections. We assessed classifier accuracy at each site (Site name, with land use marked in brackets: FOR = forest, DEV = developed site) via visual inspection with a threshold of ≤15% false positives. We applied the same species-specific classifiers to each site, to prevent site-specific differences in base classifier performance (though given the challenge of applying a single classifier across 24 sites, there were several cases where classifiers did not meet our accuracy threshold (0s in columns 2-4). We achieved accurate classifiers at a range of sites for three species: the large-billed crow (*Corvus macrorhynchos*); the Japanese bush warbler (*Horornis diphone*); and the Ryukyu scops-owl (*Otus elegans*). Accurate classifiers are marked with 1, indicating that we used data from that site-by-species combination for analysis (see *Analyses on automated species detections*). Note that *O. elegans* is a forest specialist, so is not expected to be found at the developed sites (we did not produce an accurate classifier at any developed sites for this species, in accordance with expectation). Classifiers were produced by S.R.P-J.R. and R.M. and accuracy was visually checked by a single observer (S.R.P-J.R.) for consistency.

| Site name (Land use) | *Corvus macrorhynchos* | *Horornis diphone* | *Otus elegans* |
|---:|:---:|:---:|:---:|
| Genka (FOR) | 1 | 0 | 0 |
| Hentona (FOR) | 1 | 1 | 1 |
| Katsuudake (FOR) | 1 | 1 | 1 |
| Kemin (FOR) | 1 | 1 | 1 |
| Manabi (FOR) | 1 | 1 | 0 |
| OIST forest (FOR) | 1 | 1 | 1 |
| OIST campus (FOR) | 1 | 1 | 0 |
| Oku (FOR) | 1 | 1 | 1 |
| Takeyanbaru (FOR) | 1 | 1 | 1 |
| Yona (FOR) | 1 | 1 | 1 |
| Chatan (DEV) | 0 | 0 | - |
| Gesashi (DEV) | 1 | 0 | - |
| Heiwa (DEV) | 0 | 1 | - |
| Kurashiki (DEV) | 1 | 1 | - |
| Nago (DEV) | 1 | 0 | - |
| Nakagusuku (DEV) | 1 | 1 | - |
| Oyama (DEV) | 0 | 0 | - |
| Sefa-utaki (DEV) | 1 | 1 | - |
| Senbaru (DEV) | 1 | 1 | - |
| Sueyoshi (DEV) | 1 | 0 | - |
| Tamagusuku (DEV) | 1 | 1 | - |
| Tounan (DEV) | 1 | 1 | - |
| Uehara (DEV) | 1 | 0 | - |
| Yacho (DEV) | 1 | 1 | - |
| **Total** | 21 | 17 | 7 |

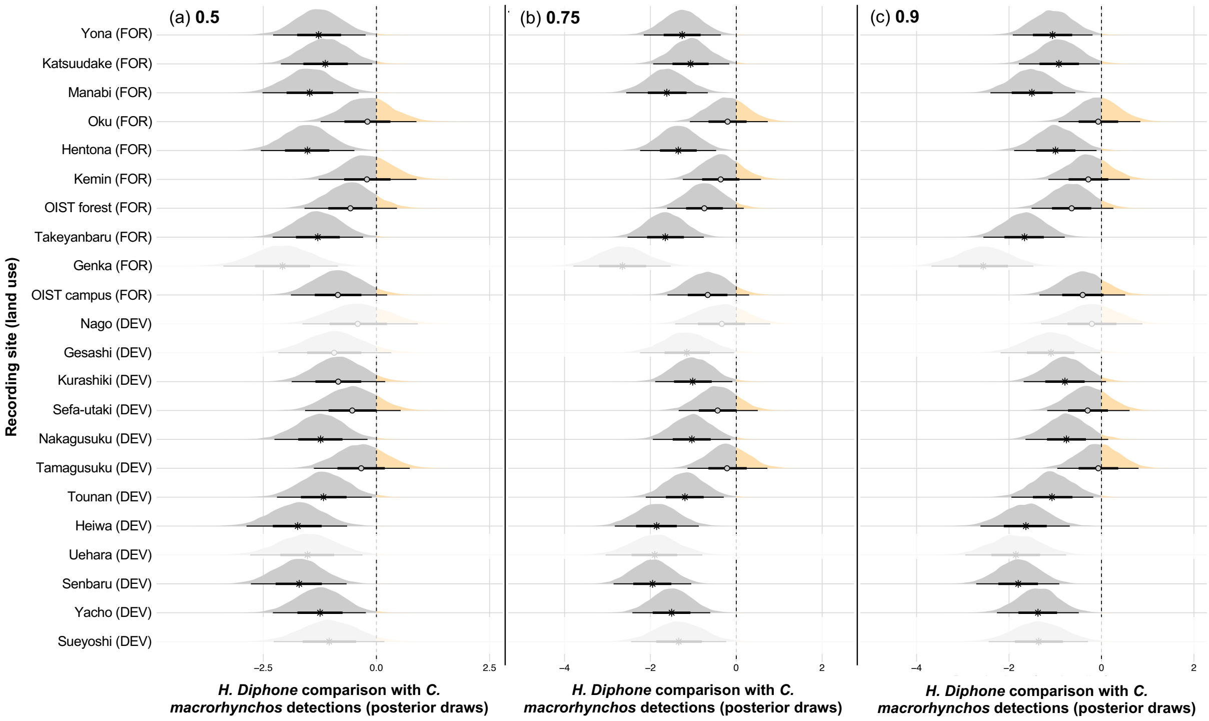


**Figure S3. Pre- and post-typhoon difference in mean detections of *Horornis diphone* and *Corvus macrorhynchos* under different detection confidence thresholds**. Posterior distributions represent 90,000 post-convergence MCMC draws of the comparison between *Horornis diphone* and *Corvus macrorhynchos* mean daily detections, where values below zero (grey) indicate lower number of detections, and values above zero (blue) a more detections of *H. diphone* relative to *C. macrorhynchos*. Non-zero-spanning credible intervals are marked with *, while circles indicate zero-spanning credible intervals (no change based on the posterior distribution). Draws are shown per site, ordered from most forested (top) to most developed (bottom) based on principle component axis 1 of the land use dimensionality reduction (Fig. S1). Panels represent results for under three automated detection probability thresholds: 0.5 (a), 0.75 (b), and 0.9 (c). Note that 0.5, which we use throughout, is a conservative filter, as credible intervals more often span zero. Inferred posterior draws (automatically computed through the site random effect term) extrapolated to field sites where species were not present (Table S1) are shown as faded distributions.

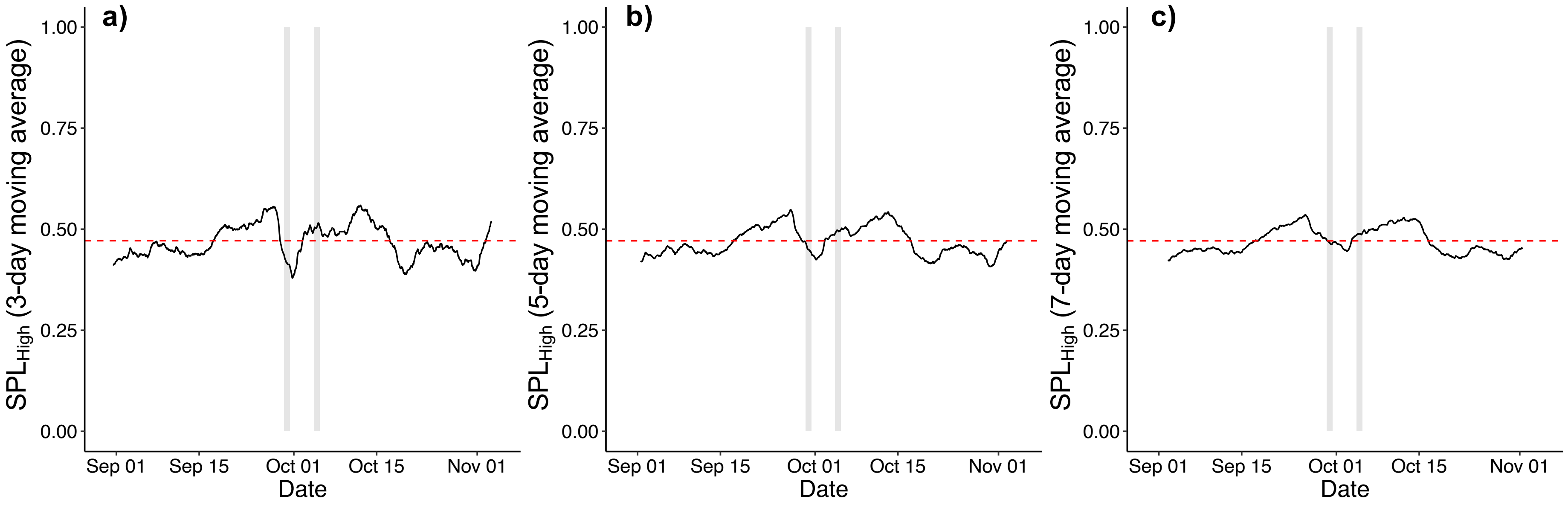


**Figure S4.** Comparison of 2-11 kHz Sound Pressure Level ($SPL_{High}$) time series at the Manabi (FOR) field site after detrending using moving average window sizes of **a**) 3 days, **b**) 5 days, and **c**) 7 days. Red dashed lines represent the pre-disturbance baseline value (mean). Grey windows indicate the periods of typhoons Trami (29-30 Sep 2018) and Kong-Rey (04-05 Oct 2018).

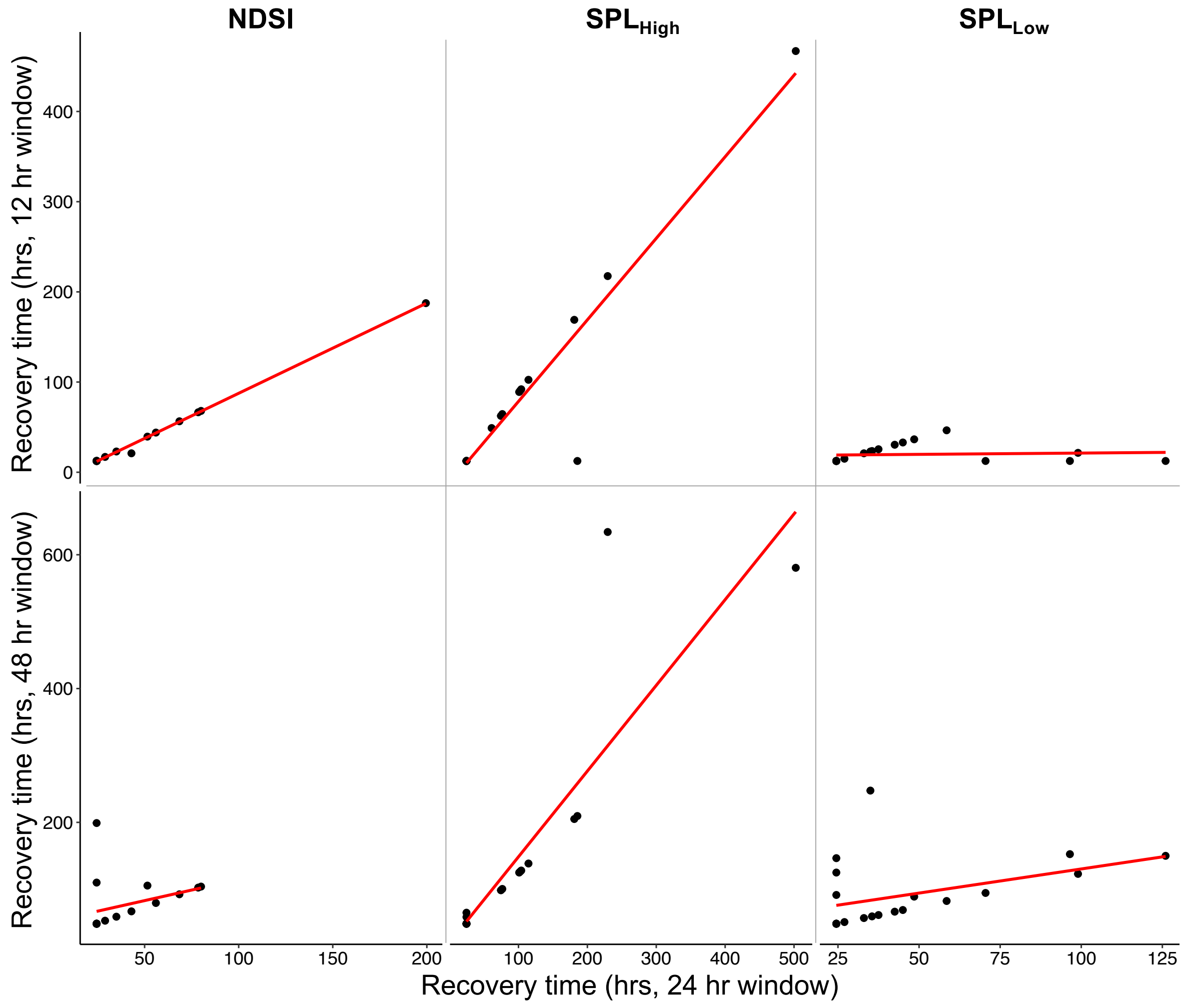


**Figure S5. Comparison of recovery time values for different recovery window sizes**. A change in window size represents a change in the amount of time for which consecutive data points must remain within the pre-typhoon baseline mean ± 1 standard deviation. Relationships between the 24 hr window size used in our analyses (x-axis) and the 12 (top panels) or 48 hr window sizes (bottom panels) are shown for three acoustic indices: NDSI (left), $SPL_{High}$, and $SPL_{Low}$ (right). Note the number of data points differs particularly for the 48 hr window size, since normalised acoustic index values were found not to recover within 30 days of the typhoon when using the 48 hr window size. Results generally show a positive correlation between the chosen 24 hr window size and other window sizes, excepting $SPL_{Low}$ with a 12 hr window size, which recovered quickly in most cases, resulting in a flat relationship. See Table S4 for results of land cover effects on recovery time for each acoustic index.

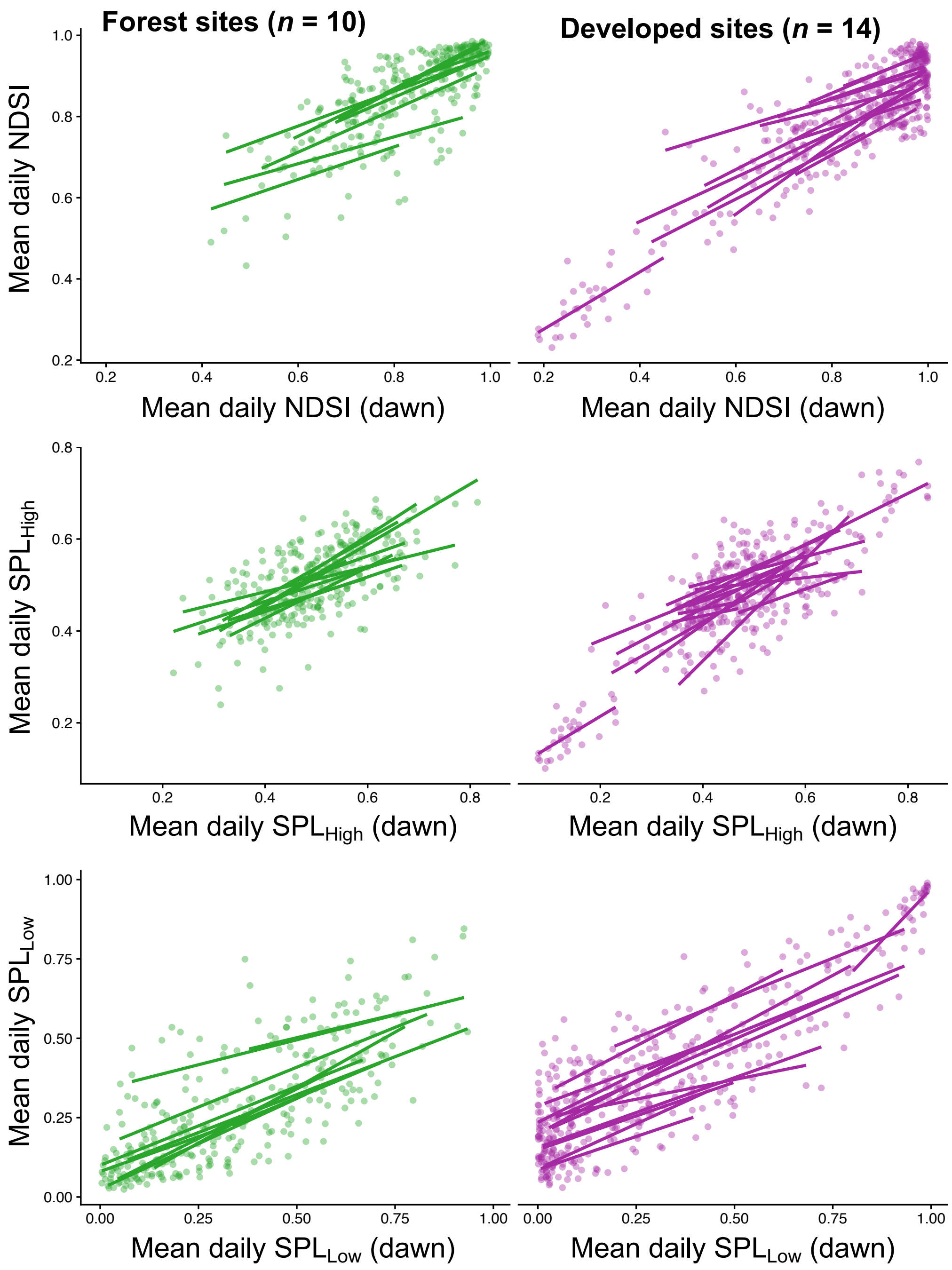


**Figure S6. Relationships between dawn sampling and daily sampling of acoustic indices**. Dawn chorus time series (x-axes) were averaged as means per day between 05:00 and 06:30. Daily time series (y-axes) were averaged as means per day across all times. The consistent positive relationship between these means suggests the daily time series used throughout our analyses was a decent approximation of dawn-only time series (another approach to isolating ecologically meaningful acoustic data). Data are separated by land use category based on *k*-means clustering (Figure S1), with the 10 forest field sites on the left (green) and 14 developed field sites on the right (purple). Y-axis values represent three acoustic indices: the Normalised Difference Soundscape Index (top), 2-11 kHz sound pressure levels [$SPL_{High}$] (middle), and 1-2 kHz sound pressure levels [$SPL_{Low}$] (bottom).

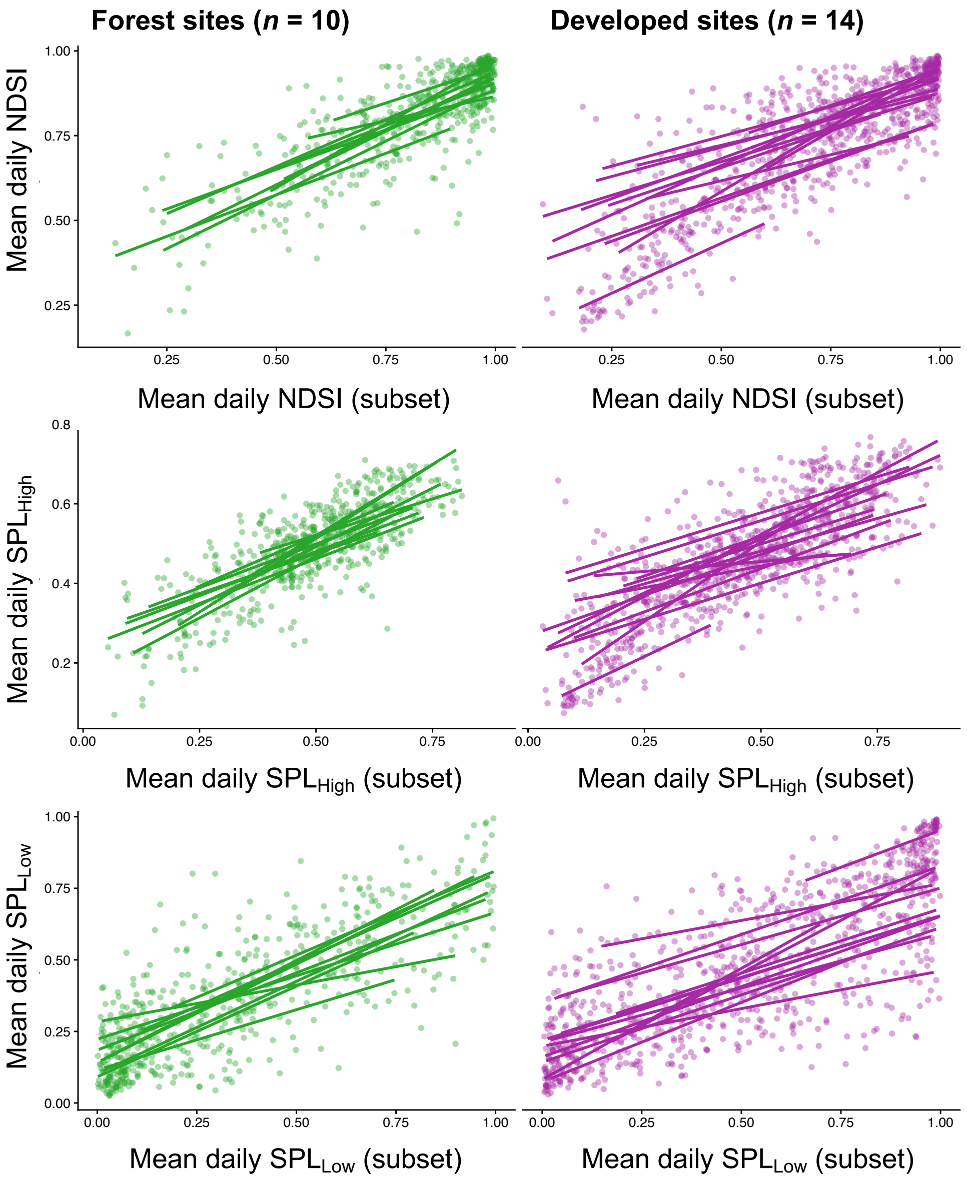


**Figure S7. Relationships between temporal subset sampling and daily sampling of acoustic indices**. Temporal subsets (x-axes) were averaged as means per day between 11:00-12:00, 14:00-15:00, and 21:00-22:00. Daily time series (y-axes) were averaged as means per day across all times. The consistent positive relationship between these means suggests the daily time series used throughout our analyses was a decent approximation of a non-dawn temporal subset of the time series (another approach to isolating ecologically meaningful acoustic data). Data are separated by land use category based on *k*-means clustering (Figure S1), with the 10 forest field sites on the left (green) and 14 developed field sites on the right (purple). Y-axis values represent three acoustic indices: the Normalised Difference Soundscape Index (top), 2-11 kHz sound pressure levels [$SPL_{High}$] (middle), and 1-2 kHz sound pressure levels [$SPL_{Low}$] (bottom).

**Table S3**. Response variables measured in this study, methods of their measurement and interpretation. Response variables were calculated separately for time series of acoustic indices and bird species detections, based largely on methods adapted from Hillebrand *et al.* (2018) and White *et al.* (2020). Following calculation, all stability measures were normalised to 0-1 (see Methods) and then temporal variability and recovery time were taken as 1 — values. This was done for ease of interpretation; all stability measures have high values which represent higher stability.

| Response variable | Time window of measurement | Method of measurement: *acoustic indices* | Method of measurement: *species detections* | Interpretation |
|---|---|---|---|---|
| **Mean state** | 30 days pre- and post-typhoon | Mean acoustic index value across the 30-day detrended time series before or after the typhoons. | Mean number of daily species detections across the 30-day time series before or after the typhoons. | Higher values correspond to higher acoustic index values (a proxy for biodiversity for NDSI and $SPL_{High}$ or higher abiotic/anthropogenic noise for $SPL_{Low}$) or more species detections. |
| **Temporal stability** | 30 days pre- and post-typhoon | 1 — coefficient of variation (that is, standard deviation/mean) among acoustic index values across the 30-day detrended time series before or after the typhoons. | 1 — coefficient of variation (that is, standard deviation / mean) among daily species detections across the 30-day time series before or after the typhoons. | Higher values correspond to lower variability through time (higher stability). |
| **Resistance** | 2 days post-typhoon | Maximum absolute difference in acoustic index values from pre-typhoon mean in the 48 hours following the 2nd typhoon. | - | Values represent log response ratios, where zero equates to complete resistance (no change), and more extreme positive or negative values represent lower resistance through over- or underperformance, respectively (Hillebrand et al., 2018). |
| **Recovery time** | 30 days post-typhoon | 1 – *recovery time*, calculated as the time taken (in hours) for acoustic index values to return to baseline (that is, mean ± 95% confidence interval of 30-day pre-typhoon detrended time series) and stay within this range for 24 consecutive hours, starting from the point of maximum displacement (the resistance point; (Garnier et al., 2017; White et al., 2020). | - | Higher values indicate shorter recovery time (higher stability). |
| **Spatial variability** | One value per recording (indices) or day (Species) across time series | Coefficient of variation (standard deviation/mean) among field sites in half-hourly mean acoustic index values (Donohue et al., 2013). | Coefficient of variation among field sites in daily species detections on each date. | Higher values represent higher variability in space, which is stabilising since spatial variability represents asynchronous biomass fluxes within or among species, in turn providing spatial insurance through patch dynamics (Leibold et al., 2004; Loreau et al., 2003). |

**Table S4**. **Mean, standard deviation (S.D.), and 95% credible intervals of the posterior distribution for parameters included in each best-fitting model.** Error distribution represents the best performing model error distribution for each variable, and the R function within the brms package used to assign each distribution. Only best performing models are reported (model performance was compared using LOOIC).

| Data | Response variable | Error distribution | Model Parameter | Mean | S.D. | 2.5% C.I. | 97.5% C.I. |
|---|---|---|---|---|---|---|---|
| Acoustic Indices | | | | | | | |
| NDSI | **Mean** | Beta | **Intercept** | **1.45** | **0.23** | **0.99** | **1.92** |
| | | | $Land_{Dev}$ | 0.37 | 0.36 | -0.35 | 1.09 |
| | | | **$Typhoon_{Post}$** | **-0.52** | **0.13** | **-0.78** | **-0.26** |
| | | | $Land_{Dev}$:$Typhoon_{Post}$ | 0.06 | 0.21 | -0.36 | 0.48 |
| | Temporal Stability | Beta | Intercept | -0.16 | 0.39 | -0.93 | 0.62 |
| | | | $Land_{Dev}$ | 0.36 | 0.59 | -0.80 | 1.53 |
| | | | $Typhoon_{Post}$ | -0.10 | 0.40 | -0.91 | 0.69 |
| | | | $Land_{Dev}$:$Typhoon_{Post}$ | -0.22 | 0.80 | -1.82 | 1.38 |
| | Resistance | Beta | Intercept | 0.66 | 0.48 | -0.29 | 1.63 |
| | | | $Land_{Dev}$ | 0.04 | 0.77 | -1.51 | 1.55 |
| | Recovery Time | Beta | Intercept | 0.94 | 0.52 | -0.08 | 1.97 |
| | | | $Land_{Dev}$ | 0.66 | 0.89 | -1.10 | 2.43 |
| $SPL_{High}$ | Mean | Beta | Intercept | -0.12 | 0.14 | -0.39 | 0.16 |
| | | | $Land_{Dev}$ | 0.16 | 0.22 | -0.26 | 0.59 |
| | | | $Typhoon_{Post}$ | -0.06 | 0.13 | -0.31 | 0.19 |
| | | | $Land_{Dev}$:$Typhoon_{Post}$ | -0.02 | 0.20 | -0.41 | 0.37 |
| | Temporal Stability | Beta | Intercept | -0.33 | 0.41 | -1.15 | 0.49 |
| | | | $Land_{Dev}$ | 0.26 | 0.61 | -0.96 | 1.46 |
| | | | $Typhoon_{Post}$ | 0.40 | 0.54 | -0.67 | 1.48 |
| | | | $Land_{Dev}$:$Typhoon_{Post}$ | -0.10 | 1.00 | -2.09 | 1.89 |
| | Resistance | Beta | Intercept | 0.10 | 0.55 | -0.98 | 1.20 |
| | | | $Land_{Dev}$ | 0.57 | 0.89 | -1.17 | 2.34 |
| | Recovery Time | Beta | Intercept | 0.91 | 0.59 | -0.26 | 2.08 |
| | | | $Land_{Dev}$ | 1.62 | 0.96 | -0.25 | 3.54 |

| Data | Response variable | Error distribution | Model Parameter | Mean | S.D. | 2.5% C.I. | 97.5% C.I. |
|---|---|---|---|---|---|---|---|
| $SPL_{Low}$ | **Mean** | Beta | Intercept | -0.51 | 0.26 | -1.01 | 0.00 |
| | | | $Land_{Dev}$ | -0.44 | 0.40 | -1.23 | 0.34 |
| | | | **$Typhoon_{Post}$** | **0.68** | **0.17** | **0.34** | **1.00** |
| | | | $Land_{Dev}$:$Typhoon_{Post}$ | -0.12 | 0.26 | -0.62 | 0.40 |
| | Temporal Stability | Beta | Intercept | 0.48 | 0.28 | -0.07 | 1.04 |
| | | | $Land_{Dev}$ | -0.98 | 0.72 | -2.42 | 0.46 |
| | | | $Typhoon_{Post}$ | -0.73 | 0.45 | -1.62 | 0.16 |
| | | | $Land_{Dev}$:$Typhoon_{Post}$ | 0.34 | 0.95 | -1.55 | 2.24 |
| | Resistance | Beta | Intercept | 0.20 | 0.48 | -0.73 | 1.15 |
| | | | $Land_{Dev}$ | 0.30 | 0.77 | -1.21 | 1.84 |
| | Recovery Time | Beta | Intercept | 0.04 | 0.53 | -1.01 | 1.07 |
| | | | $Land_{Dev}$ | 1.11 | 0.64 | -0.15 | 2.40 |
| <u>Species detections</u> | | | | | | | |
| | **Mean Daily Detections** | lognormal | **Intercept** | **3.57** | **0.32** | **2.95** | **4.21** |
| | | | $Land_{Dev}$ | 0.25 | 0.45 | -0.64 | 1.14 |
| | | | $Typhoon_{Post}$ | -0.17 | 0.35 | -0.86 | 0.51 |
| | | | **$Species_{Horo}$** | **-1.05** | **0.39** | **-1.82** | **-0.28** |
| | | | $Species_{Otus}$ | -0.47 | 1.43 | -3.29 | 2.34 |
| | | | $Land_{Dev}$:$Typhoon_{Post}$ | -0.43 | 0.49 | -1.39 | 0.53 |
| | | | $Land_{Dev}$:$Species_{Horo}$ | -0.35 | 0.53 | -1.38 | 0.70 |
| | | | $Land_{Dev}$:$Species_{Otus}$ | -0.47 | 1.43 | -3.27 | 2.34 |
| | | | **$Typhoon_{Post}$:$Species_{Horo}$** | **-1.27** | **0.52** | **-2.29** | **-0.24** |
| | | | $Typhoon_{Post}$:$Species_{Otus}$ | -0.06 | 1.45 | -2.91 | 2.79 |
| | | | $Land_{Dev}$:$Typhoon_{Post}$:$Species_{Horo}$ | 0.15 | 0.71 | -1.25 | 1.53 |
| | | | $Land_{Dev}$:$Typhoon_{Post}$:$Species_{Otus}$ | -0.07 | 1.45 | -2.91 | 2.77 |

| Data | Response variable | Error distribution | Model Parameter | Mean | S.D. | 2.5% C.I. | 97.5% C.I. |
|---|---|---|---|---|---|---|---|
| | **Temporal Stability** | lognormal | **Intercept** | **0.34** | **0.05** | **0.25** | **0.43** |
| | | | $Land_{Dev}$ | 0.00 | 0.07 | -0.12 | 0.14 |
| | | | **$Typhoon_{Post}$** | **0.23** | **0.05** | **0.13** | **0.33** |
| | | | $Species_{Horo}$ | 0.07 | 0.06 | -0.04 | 0.19 |
| | | | $Species_{Otus}$ | -0.09 | 1.41 | -2.86 | 2.68 |
| | | | $Land_{Dev}$:$Typhoon_{Post}$ | -0.09 | 0.07 | -0.24 | 0.06 |
| | | | $Land_{Dev}$:$Species_{Horo}$ | 0.04 | 0.08 | -0.12 | 0.20 |
| | | | $Land_{Dev}$:$Species_{Otus}$ | -0.07 | 1.41 | -2.83 | 2.70 |
| | | | $Typhoon_{Post}$:$Species_{Horo}$ | -0.11 | 0.08 | -0.27 | 0.05 |
| | | | $Typhoon_{Post}$:$Species_{Otus}$ | -0.01 | 1.42 | -2.81 | 2.77 |
| | | | $Land_{Dev}$:$Typhoon_{Post}$:$Species_{Horo}$ | -0.04 | 0.11 | -0.26 | 0.18 |
| | | | $Land_{Dev}$:$Typhoon_{Post}$:$Species_{Otus}$ | -0.03 | 1.42 | -2.80 | 2.77 |

**Table S5. Mean and 95% credible intervals of the posterior distribution for intercept segments included in each best-fitting break-point model.** Site subset reports the subset of the data over which spatial variability was calculated (All = all 24 sites, Forest = 10 forest sites, Developed = 14 developed sites). The time agreement of the break points is reported (after setting weakly informative priors, see main text) as either strong (all break points show little-no variation in the timing of the breaks), medium (all break points show a little variation, or a single break point shows some variation in the timing of breaks), or weak (break points have large variation in the timing of breaks). We also report whether break points are coincident with the typhoons (after setting weakly informative priors), where "yes" indicates that one or several break points closely align with the timing of the typhoons. Segment indicates the time period captured by each intercept segment (i.e., a period before a break point), and contrasts indicate pairwise contrasts based on nonoverlapping 95% credible intervals; where A is different from B but not A or AB ($p$ = 0.05 equivalent), for instance. Though we used weakly informative priors as starting points for the break point models, model selection via LOOIC can favour models without break points if no such points exist. Only best performing models are reported (based on LOOIC). Note: for the forest specialist, Otus elegans, "All" is equivalent to the forest site subset, so we report only the latter here. See Figures S8 and S12 for break-point model outputs for the spatial variability of acoustic indices and species detections, respectively.

| Data | Site subset | Break points | Time agreement? | Typhoon coincident? | Segment | Mean | 2.5% C.I. | 97.5% C.I. | contrasts |
|---|---|---|---|---|---|---|---|---|---|
| <u>Acoustic Indices</u> | | | | | | | | | |
| NDSI | All | 3 | Strong | Yes | Pre-typhoon | 0.171 | 0.170 | 0.172 | A |
| | | | | | Typhoon period | 0.193 | 0.186 | 0.199 | B |
| | | | | | Post-typhoon 1 | 0.211 | 0.209 | 0.214 | C |
| | | | | | Post-typhoon 2 | 0.255 | 0.253 | 0.256 | D |
| | Forest | 3 | Medium | Yes | Pre-typhoon | 0.112 | 0.108 | 0.118 | A |
| | | | | | Typhoon period | 0.147 | 0.094 | 0.169 | AB |
| | | | | | Post-typhoon 1 | 0.114 | 0.105 | 0.134 | A |
| | | | | | Post-typhoon 2 | 0.172 | 0.168 | 0.176 | B |
| | Developed | 3 | Strong | Yes | Pre-typhoon | 0.201 | 0.200 | 0.203 | A |
| | | | | | Typhoon period | 0.219 | 0.214 | 0.226 | B |
| | | | | | Post-typhoon 1 | 0.249 | 0.245 | 0.252 | C |
| | | | | | Post-typhoon 2 | 0.294 | 0.291 | 0.296 | D |

| Data | Site subset | Break points | Time agreement? | Typhoon coincident? | Segment | Mean | 2.5% C.I. | 97.5% C.I. | contrasts |
|---|---|---|---|---|---|---|---|---|---|
| $SPL_{High}$ | All | 3 | Strong | Yes | Pre-typhoon | 0.194 | 0.191 | 0.196 | A |
| | | | | | Typhoon period | 0.235 | 0.225 | 0.245 | B |
| | | | | | Post-typhoon 1 | 0.275 | 0.270 | 0.281 | C |
| | | | | | Post-typhoon 2 | 0.344 | 0.340 | 0.347 | D |
| | Forest | 3 | Medium/Strong | Yes | Pre-typhoon | 0.114 | 0.110 | 0.117 | A |
| | | | | | Typhoon period | 0.197 | 0.149 | 0.227 | B |
| | | | | | Post-typhoon 1 | 0.199 | 0.189 | 0.212 | B |
| | | | | | Post-typhoon 2 | 0.273 | 0.269 | 0.278 | C |
| | Developed | 3 | Strong | Yes | Pre-typhoon 1 | 0.234 | 0.229 | 0.238 | A |
| | | | | | Pre-typhoon 2 | 0.265 | 0.240 | 0.286 | B |
| | | | | | Post-typhoon 1 | 0.324 | 0.317 | 0.332 | C |
| | | | | | Post-typhoon 2 | 0.390 | 0.386 | 0.394 | D |
| $SPL_{Low}$ | All | 3 | Medium/Strong | Yes | Pre-typhoon | 0.648 | 0.631 | 0.666 | A |
| | | | | | Typhoon period | 0.371 | 0.340 | 0.401 | B |
| | | | | | Post-typhoon 1 | 0.525 | 0.497 | 0.573 | C |
| | | | | | Post-typhoon 2 | 0.381 | 0.342 | 0.454 | B |
| | Forest | 3 | Strong | Yes | Pre-typhoon | 0.635 | 0.622 | 0.648 | A |
| | | | | | Typhoon period | 0.325 | 0.303 | 0.347 | B |
| | | | | | Post-typhoon 1 | 0.428 | 0.415 | 0.442 | C |
| | | | | | Post-typhoon 2 | 0.324 | 0.302 | 0.348 | B |
| | Developed | 3 | Medium/Strong | Yes | Pre-typhoon | 0.619 | 0.600 | 0.641 | A |
| | | | | | Typhoon period | 0.390 | 0.338 | 0.418 | B |
| | | | | | Post-typhoon 1 | 0.489 | 0.467 | 0.511 | C |
| | | | | | Post-typhoon 2 | 0.363 | 0.331 | 0.396 | B |

| Data | Site subset | Break points | Time agreement? | Typhoon coincident? | Segment | Mean | 2.5% C.I. | 97.5% C.I. | contrasts |
|---|---|---|---|---|---|---|---|---|---|
| <u>Species Detections</u> | | | | | | | | | |
| *C. macrorhynchos* | All | 0 | - | No | Full series | 1.346 | 1.279 | 1.420 | - |
| | Forest | 1 | Weak/Medium | Yes | Pre-typhoon | 1.133 | 0.990 | 1.275 | A |
| | | | | | Post-typhoon | 1.517 | 1.378 | 1.366 | B |
| | Developed | 1 | Weak | No | Pre-typhoon 1 | 1.670 | 1.346 | 1.993 | A |
| | | | | | Pre-typhoon 2 | 1.256 | 1.172 | 1.338 | B |
| *H. diphone* | All | 1 | Strong | Yes | Pre-typhoon | 1.408 | 1.150 | 1.663 | A |
| | | | | | Post-typhoon | 2.318 | 2.090 | 2.535 | B |
| | Forest | 1 | Strong | Yes | Pre-typhoon | 1.147 | 0.953 | 1.330 | A |
| | | | | | Post-typhoon | 1.995 | 1.817 | 2.166 | B |
| | Developed | 1 | Weak | No | Segment 1 | 1.726 | 1.359 | 2.031 | A |
| | | | | | Segment 2 | 2.402 | 1.927 | 3.198 | A |
| *O. elegans* | Forest | 0 | - | No | Full series | 3.354 | 3.201 | 3.509 | - |

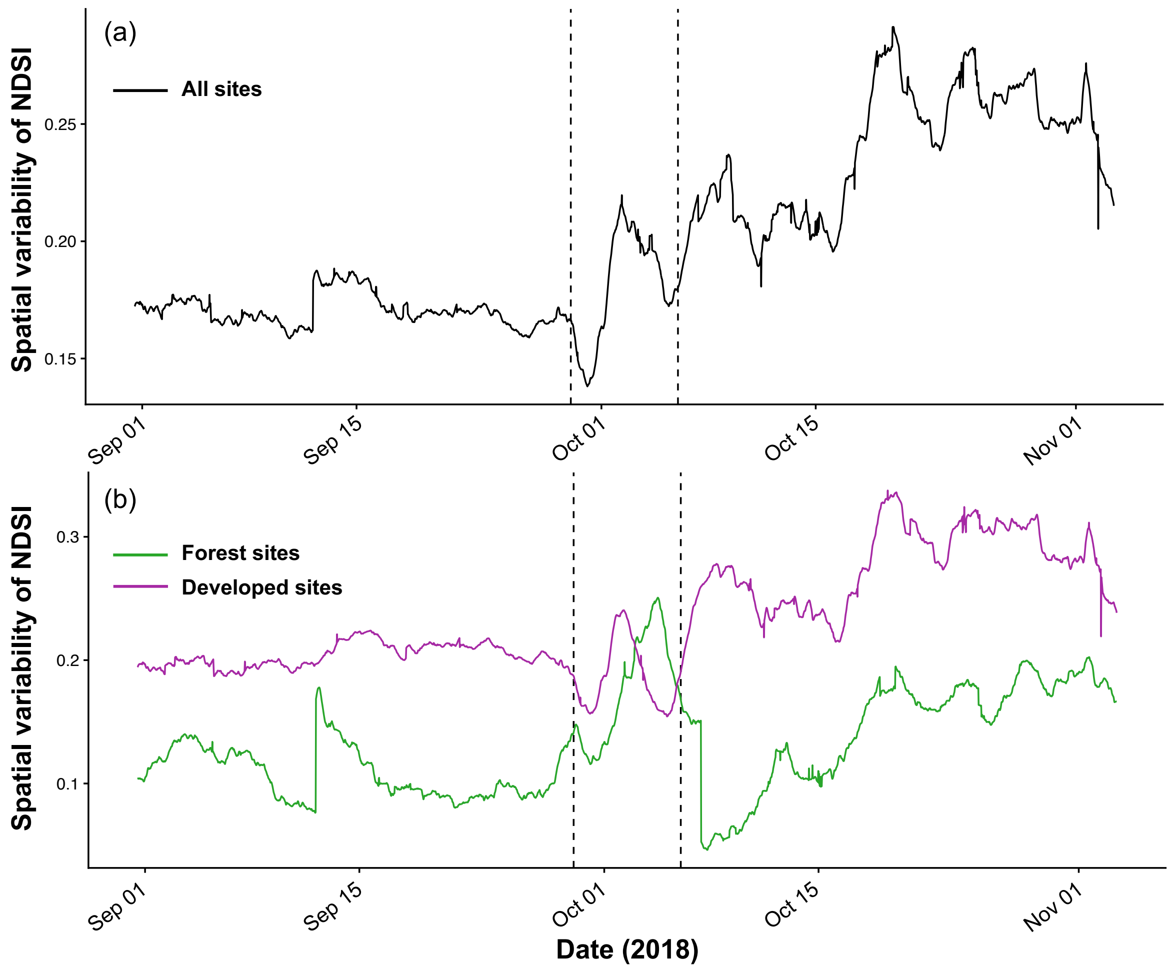


**Figure S8**. **Spatial variability of the Normalised Difference Soundscape Index [NDSI] through time**. Time series of NDSI spatial variability across all sites (a), and across forest (green) and developed (purple) sites separately (b). Dashed lines delineate the pre- and post-typhoon periods.

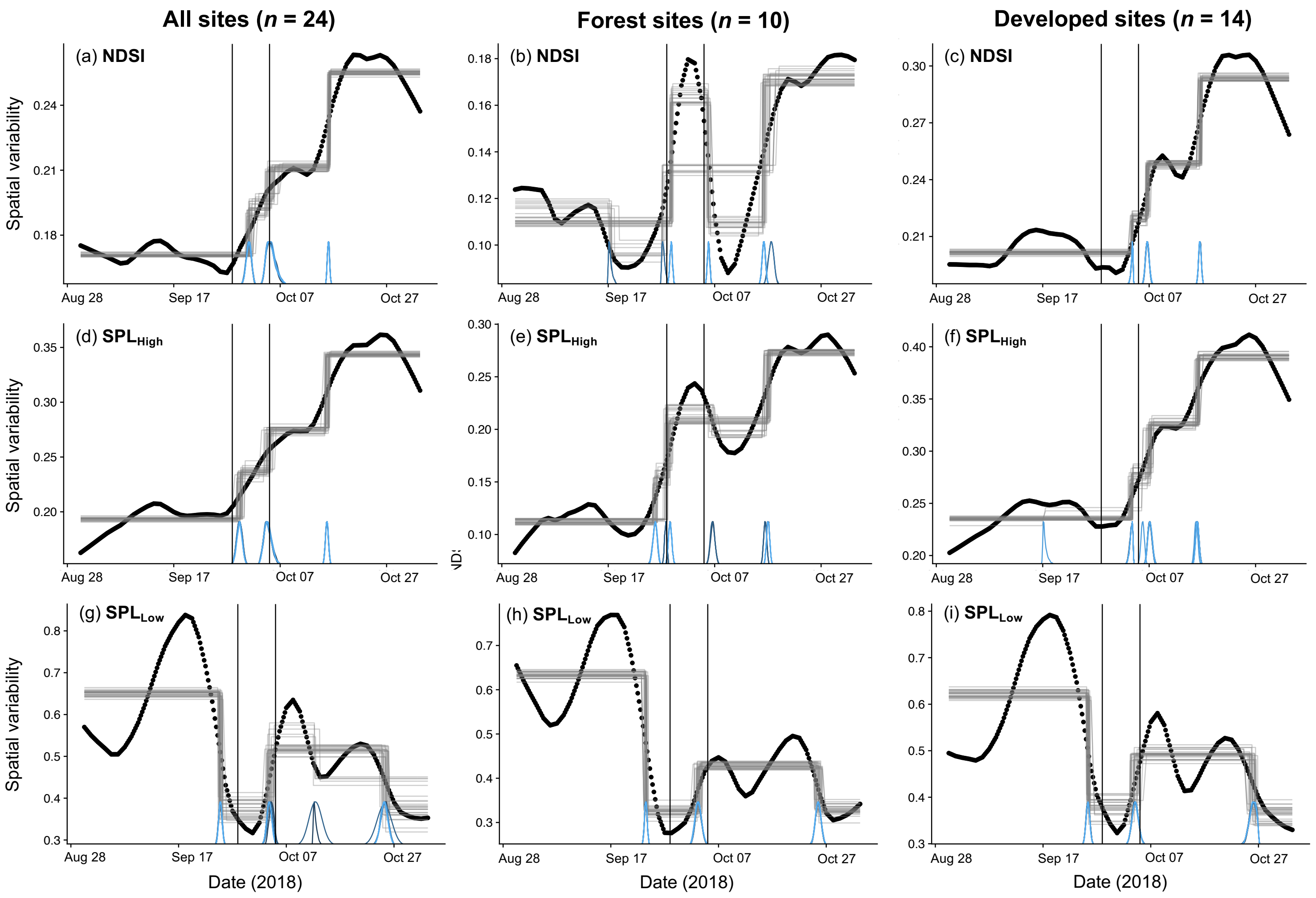


**Figure S9**. Best fitting break-point models of acoustic index spatial variability from Table S5. Spatial variability values (Y-axis) are weekly detrended values of spatial variability through time (X-axis) calculated as the coefficient of variation across all 24 sites (a,d,g), the 10 forest sites (b,e,h) or 14 developed sites (c,f,i). Acoustic indices are the normalised difference soundscape index (a-c), or its 2-11 kHz sound pressure level [$SPL_{High}$] (d-f) or 1-2 kHz sound pressure level [$SPL_{Low}$] (g-i) components. Blue frequency distributions represent posterior density distributions of the timing of break points (after weakly informative priors, see main text).

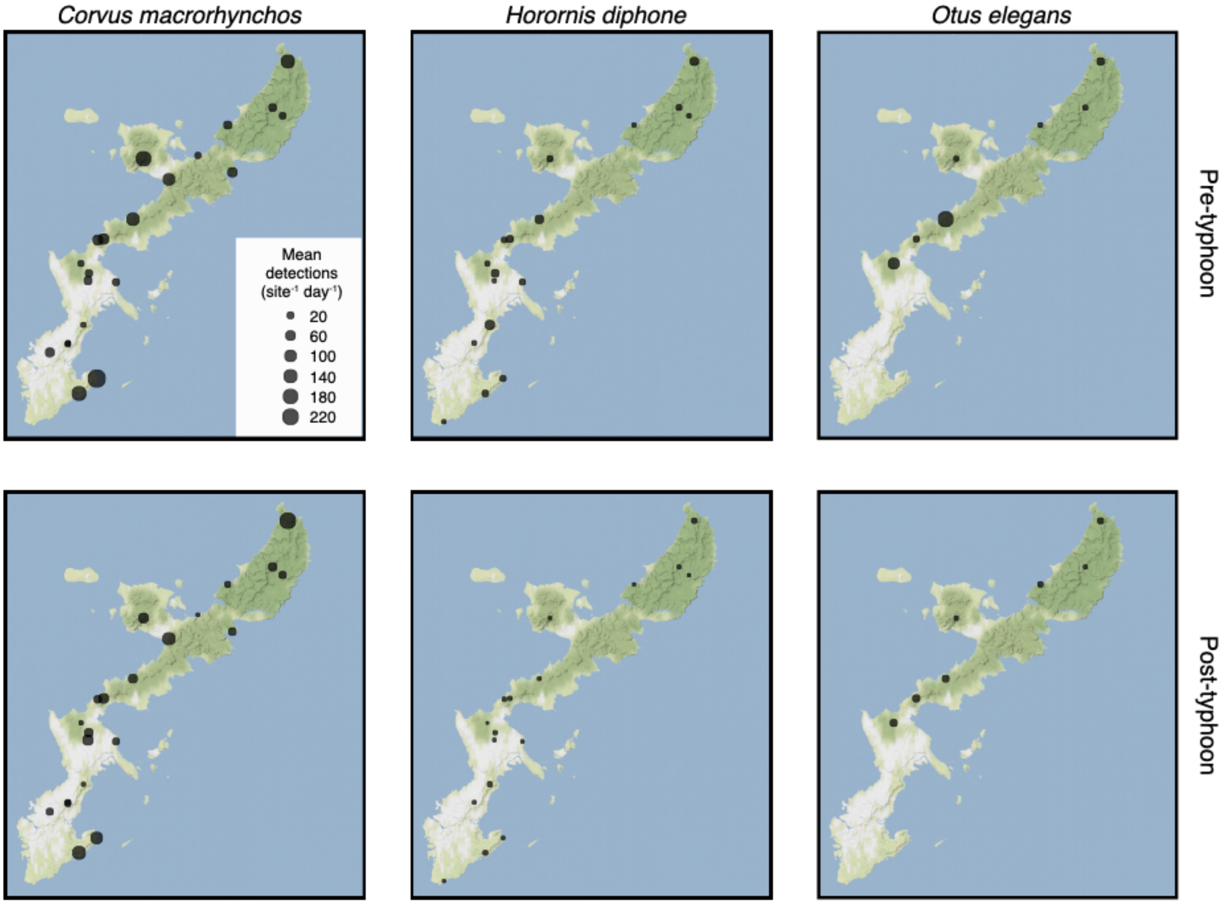


**Figure S10**. **Mean daily species detection before and after the typhoons**. Summed automated species detections for *Horornis diphone* per day at each site, averaged across the 30-day pre-typhoon period (top) and post-typhoon period (bottom) for each of our target species: the large billed crow (*Corvus macrorhynchos*, L), the Japanese bush warbler (*Horornis diphone*), and the Ryukyu scops owl (*Otus elegans*, R). Point size represents mean daily detections across each 30-day period (see legend, top left) and points are shown at the corresponding site locations on maps of Okinawa.

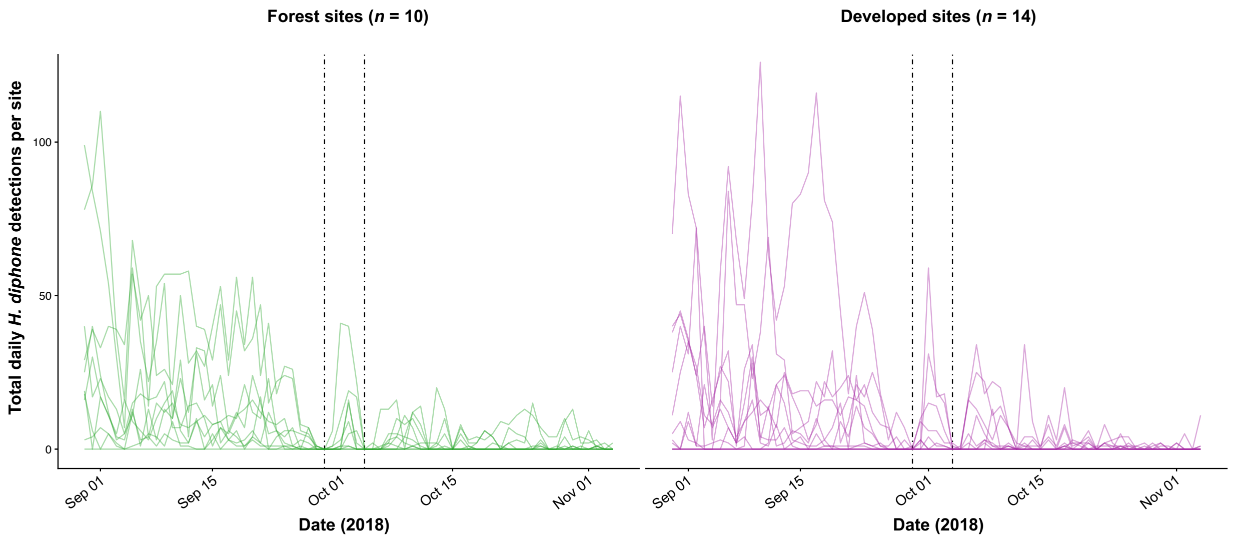


**Figure S11**. **Site-level daily detections of *Horornis diphone***. Summed automated species detections for *Horornis diphone* per day at each site for forest sites (L) and developed sites (R). Individual lines represent changes in detection at a given field site. Vertical barred lines indicate the day of each typhoon's closest pass to Okinawa. Based on the clear decline in detections during and immediately following each typhon, we infer that the observed overall post-typhoon decline in H. diphone detections (Figure 4b) was most likely related to typhoon impacts, rather than, for instance, a seasonal decline in vocalisations of this species.

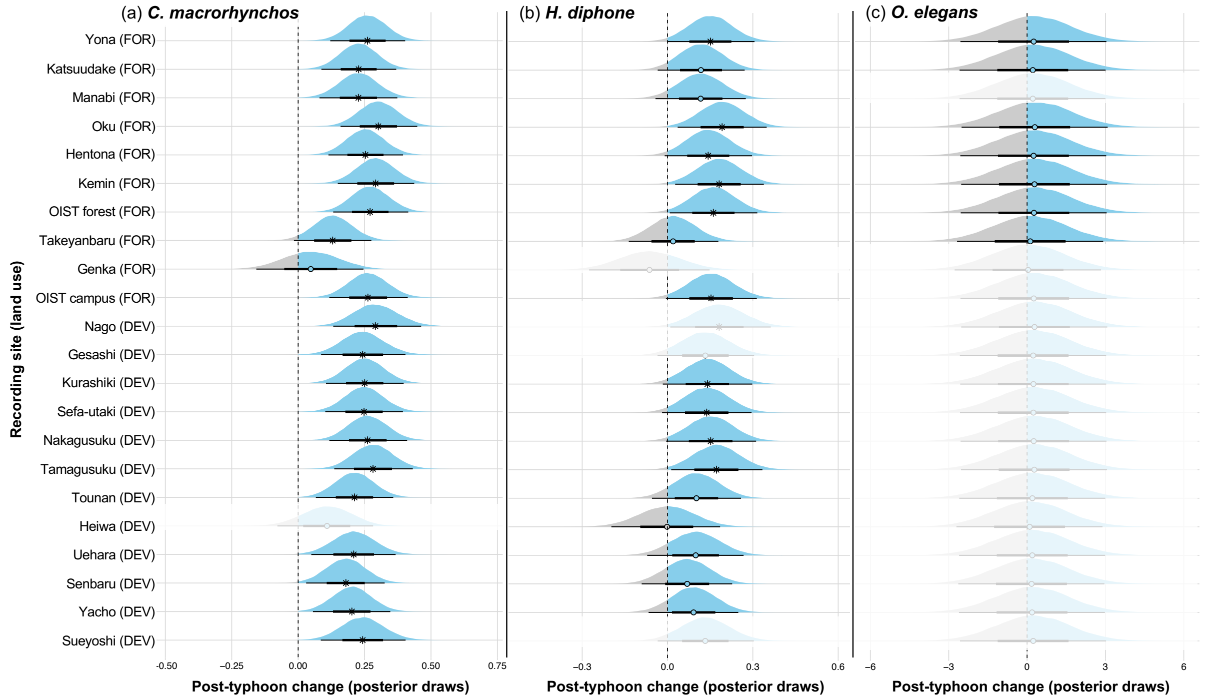


**Figure S12. Comparison of temporal stability of species detections before and after the typhoons**. Posterior distributions represent 90,000 post-convergence MCMC draws of the change from pre- to post-typhoon periods, where values below zero (grey) indicate a post-typhoon decline, and values above zero (blue) a post-typhoon increase in the temporal stability of automated species vocalisation detections, broken down by species: *Corvus macrorhynchos* (a), *Horornis diphone* (b), *Otus elegans* (c). Non-zero-spanning credible intervals are marked with *, while circles indicate zero-spanning credible intervals (no change based on the posterior distribution). Draws are shown per site, ordered from most forested (top) to most developed (bottom) based on principle component axis 1 of the land use dimensionality reduction (Fig. S1). Inferred posterior draws (automatically computed through the site random effect term) extrapolated to field sites where species were not present (Table S1) are shown as faded distributions.

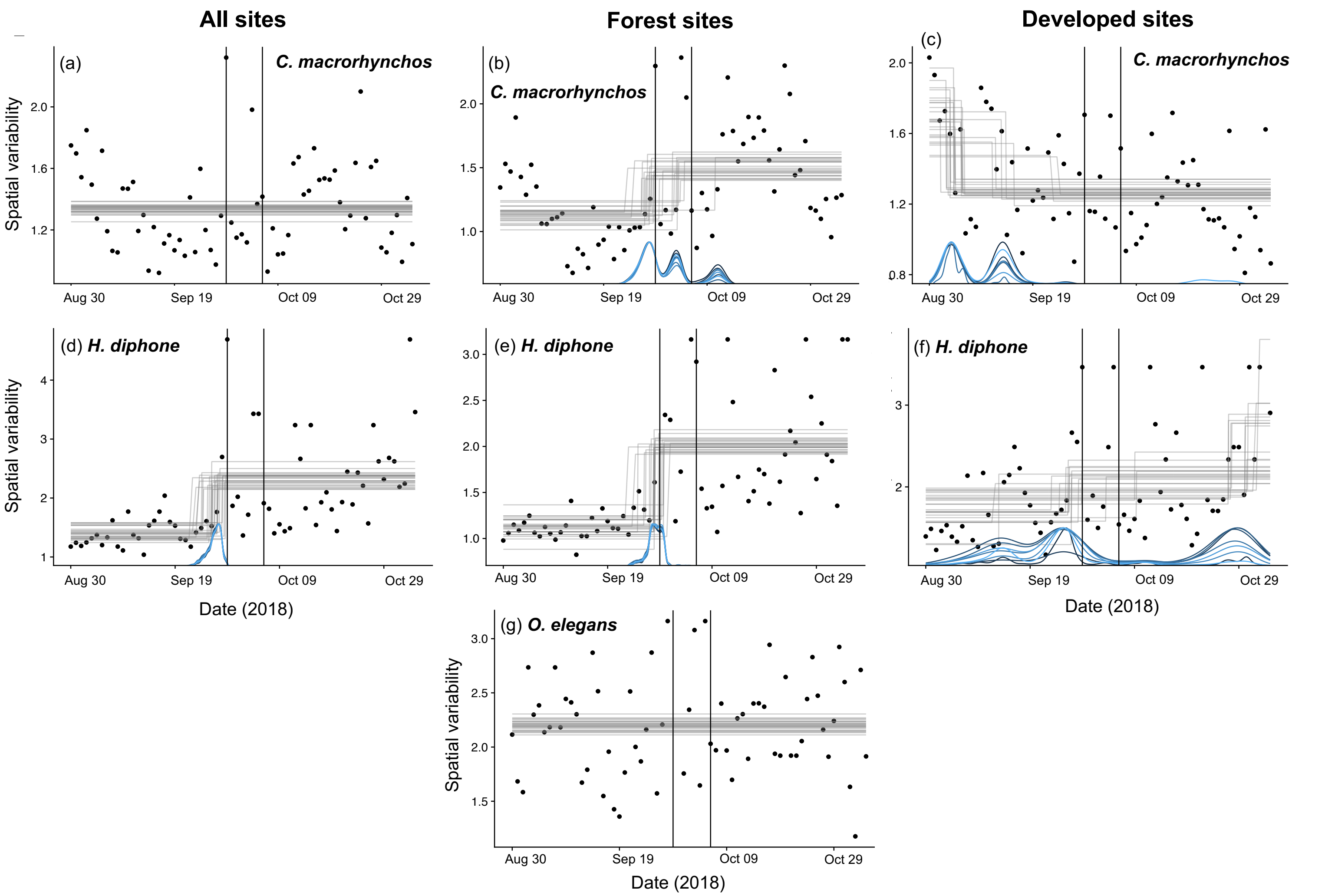


**Figure S13**. Best fitting break-point models of species detection spatial variability from Table S5. Spatial variability values (Y-axis) through time (X-axis) are calculated as the coefficient of variation across all available sites (a,d), the relevant forest sites (b,e,g) or the relevant developed sites (c,f,i). See Table S2 for the sites across which spatial variability was calculated for each species. Species are *Corvus macrorhynchos* (a-c), *Horornis diphone* (d-f) or *Otus elegans* (g). The forest specialist *O. elegans* was not detected in any developed sites. Blue frequency distributions represent posterior density distributions of the timing of break points (after weakly informative priors, see main text).
